# The mouse midbrain periaqueductal gray is organized into target-specific output populations

**DOI:** 10.64898/2026.11.06.764863

**Authors:** Alicia Strosche, Ayu Tamaki, Jonathan N. Mader, Eloah S. De Biasi, A. Vanessa Stempel

## Abstract

How brains select between behaviours and coordinate their execution is a central question in neuroscience. The vertebrate periaqueductal gray (PAG) controls a remarkable range of instinctive behaviours, from escape and freezing to hunting, vocalization, and micturition. Its columnar organization, the classical framework for mapping these functions, cannot explain how one region controls so many, often mutually exclusive, behaviours. To determine how PAG output is organized, we generated a whole-brain, column- and cell-type-specific synaptic projection atlas of the mouse PAG, showing distinct dorsal and ventral outputs and dense within-PAG connectivity. Retrograde tracing from four motor-related target regions identified segregated PAG populations, biased towards, but not confined to, individual columns. Channelrhodopsin-2-assisted circuit mapping showed that one such population, projecting to the cuneiform nucleus, provides substantial local collaterals, positioned to shape ongoing PAG activity. These findings uncover a modular output architecture that extends the columnar framework, offering a circuit logic for how one brain region produces diverse behaviours.

## Introduction

The motor behaviours by which animals interact with the world are primarily instinctive. In vertebrates, the circuits that produce them are located in evolutionarily conserved brainstem regions. Of these circuits, the periaqueductal gray (PAG) coordinates the initiation of a given behaviour as well as motor, autonomic, and antinociceptive components (1–7). The activation of sets of PAG neurons initiates complete programs of sequential movements (such as ‘escape’) rather than single motor actions (‘turn’, ‘run’, ‘stop’) (8–10). This indicates that the PAG itself contains the circuitry to assemble each program rather than passively relaying signals from elsewhere. The functional diversity of behaviours controlled by the PAG has classically been attributed to its anatomical organization into four longitudinal columns: a dorsomedial (dmPAG), dorsolateral (dlPAG) – together referred to as dorsal (dPAG) – lateral (lPAG), and ventrolateral (vlPAG) column. Each has been preferentially associated with a subset of behaviours: the dPAG with active defence, the lPAG with vocalizations, hunting, and pup retrieval, and the vlPAG with freezing, micturition, and parasympathetic responses (4, 8, 11–20). A purely columnar framework cannot, however, account for the diversity of responses evoked by PAG activation. For example, optogenetic activation of a single column can drive behaviours as distinct as freezing and micturition (16, 18), and individual columns project to many downstream targets (2, 19, 21–23). Studies targeting projection-defined populations suggest that behavioural specificity may instead arise at the level of individual output pathways. For example, glutamatergic neurons projecting to the magnocellular reticular nucleus (Mc) of the medulla elicit freezing (16), whereas glutamatergic neurons projecting to the cuneiform nucleus (CnF) elicit escape (24). Each of these pathways was characterized in isolation, leaving unclear whether they arise from genuinely dedicated neurons or from shared, or partially overlapping, populations with branching collaterals (that might, for example, carry an initiation signal).

A modular organization principle – projection-defined cell populations that project to a single specific downstream effector – is a recurring motif in circuits for motor control (25–32). The advantage of such an architecture is that a single region can issue functionally distinct, sometimes opposing, commands: a dedicated population can arrest a behaviour without disrupting other components (25), can be recruited in a specific behavioural context (26), or can suppress motor programs incompatible with the one being executed (30). In each case, the functional unit is defined by projection target rather than by anatomical position. This principle has been suggested for some PAG projections to the medulla (33). Whether this extends as a general principle for PAG output is not known (6). We therefore ask whether PAG output follows a systematic logic, and whether the many behaviours driven by the PAG are generated by target-specific subpopulations organized as modular output pathways.

We first mapped the brain-wide projections of glutamatergic (vesicular glutamate transporter 2-positive, VGluT2^+^) and GABAergic (vesicular GABA transporter-positive, VGAT^+^) PAG neurons using column- and cell-type-specific anterograde tracing at synaptic resolution. We then used dual-colour retrograde tracing to test, at single-neuron resolution, whether four motor-related target regions are innervated by overlapping or segregated PAG populations. Finally, we used channelrhodopsin-2 (ChR2)-assisted circuit mapping (CRACM) *ex vivo* to characterize the local and long-range synaptic outputs of a projection-defined PAG population, providing a circuit-level view of how a single target region combines excitation and inhibition to shape its output.

## Results

We performed small, targeted injections of a Cre-dependent virus expressing fluorescently labelled synaptophysin in the different columns of the PAG in VGluT2-Cre and VGAT-Cre mice (Methods; Figure 1A-E and 2A). The brains were aligned and registered to the Allen Mouse Brain Common Coordinate Framework version 3 (CCFv3) (34) and injection sites were hierarchically clustered based on their dorsoventral position and assigned to the PAG columns accordingly (Figure S1A-C). Given that the borders between PAG columns are not definitively defined in the literature, and that the precision of stereotaxic targeting is limited, we did not distinguish dmPAG and dlPAG in the anterograde tracing. Hierarchical clustering of the projection patterns reliably grouped dPAG injections, but did not differentiate between lateral and ventrolateral injections (Figure S1D), which are known to share broadly overlapping projection targets (4, 22), but could, in part, be attributed to the overlap of injection site locations. We therefore grouped lateral and ventrolateral injections, referred to throughout as ventral PAG (vPAG).

**Figure 1.**
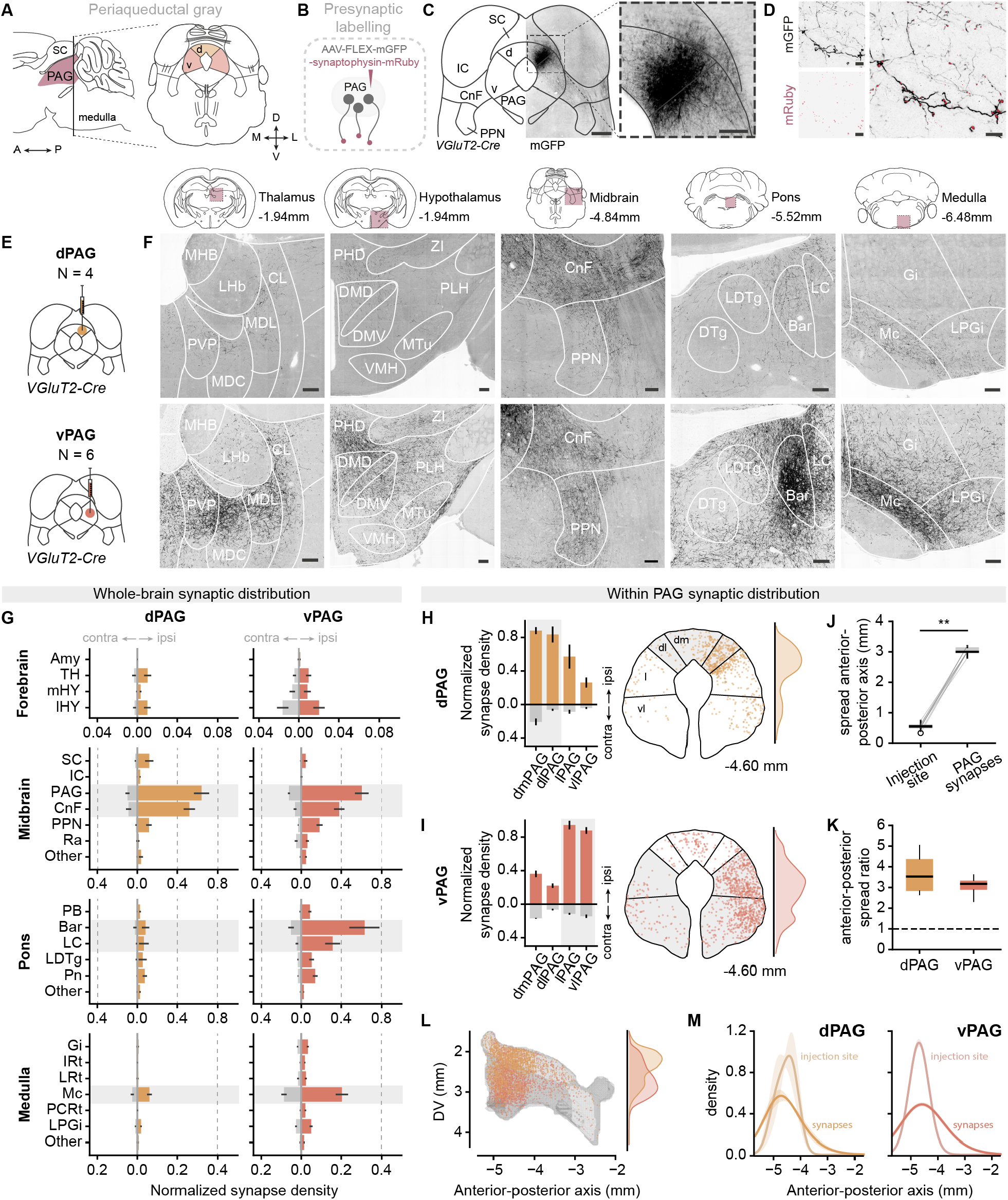
Whole-brain distribution of synaptic terminals from VGluT2^+^ PAG neurons. **(A)** Schematic of the PAG in sagittal view, indicating anterior-posterior axis (left) and in coronal view, indicating dorsoventral and mediolateral axes, and the division of the PAG into dorsal and ventral columns (right). **(B)** Schematic of the experimental design for anterograde labelling of presynaptic boutons. **(C)** Expression pattern of membrane-bound GFP (mGFP) in VGluT2^+^ dPAG neurons at the injection site (scale bars, left panel: 500 µm, right panel: 200 µm). **(D)** Expression of mGFP (black) in neuronal processes and mRuby-labelled presynaptic boutons (red) of VGluT2^+^ dPAG neurons (left panels), with an overlay of the two channels (right panel; scale bars: 10 µm). **(E)** Cell-type- and column-specific anterograde labelling strategy for VGluT2^+^ neurons from the dPAG and vPAG (top and bottom panel, respectively). **(F)** Example images of mGFP-labelled fibres in the thalamus, hypothalamus, mesencephalic locomotor region/midbrain, pons, and medulla (from left to right) of VGluT2^+^ neurons from the dPAG (top panels) and vPAG (bottom panels; scale bars, 100 µm). **(G)** Regional density of synapses from VGluT2^+^ dPAG (left panels, orange, *N =* 4 mice) and vPAG neurons (right panels, red, *N =* 6 mice), normalized to the densest region per brain. Grey and coloured bars represent contralateral and ipsilateral hemispheres, respectively. The PAG and CnF are strongly innervated across columns, while Bar, LC, and Mc are preferentially innervated by vPAG neurons (shaded areas). **(H)** Local density of synapses from VGluT2^+^ neurons of the dPAG, normalized to the densest region per brain and shown for the ipsi- and contralateral hemisphere, respectively (left panel). Example spatial distribution of VGluT2^+^ synapses within the PAG at one coronal level, downsampled by a factor of 10, and 2D kernel density estimation (KDE) of synapse positions (right panel). **(I)** Same as (H) for synapses from VGluT2^+^ neurons of the vPAG. **(J)** Within-PAG spread of the localization of synapses versus extent of the injection site along the anterior-posterior axis of the entire PAG, demonstrating that synaptic spread significantly exceeds the injection site (*N =* 10 mice, paired Wilcoxon signed-rank test: *p* < 0.01). **(K)** Ratio of within-PAG spread of synapses over the extent of the injection site by column of injection. The dashed line indicates a spread ratio of 1 (synaptic spread equals injection site extent). The ratio does not differ between the injection groups (Mann-Whitney U test: *p* = 0.610). **(L)** Reconstruction of synaptic distribution within the PAG from dPAG (orange) and vPAG neurons (red), downsampled by a factor of 400, and 2D KDE of synapse positions along the dorsoventral axis. **(M)** KDE of injection site area and number of synapses along the anterior-posterior extent of the PAG by injection column (dPAG: left panel, orange; vPAG: right panel, red). Amy, amygdala; Bar, Barrington’s nucleus; CL, centrolateral thalamic nucleus; CnF, cuneiform nucleus; DMD, dorsomedial hypothalamic nucleus, dorsal part; DMV, dorsomedial hypothalamic nucleus, ventral part; DTg, dorsal tegmental nucleus; Gi, gigantocellular reticular nucleus; IC, inferior colliculus; IRt, intermediate reticular nucleus; LC, locus coeruleus; LDTg, laterodorsal tegmental nucleus; LHb, lateral habenula; lHY, lateral hypothalamus; LPGi, lateral paragigantocellular nucleus; LRt, lateral reticular nucleus; MDC, mediodorsal thalamic nucleus, central; MDL, mediodorsal thalamic nucleus, lateral; MHB, medial habenular nucleus; mHY, medial hypothalamus; Mc, magnocellular reticular nucleus; MTu, medial tuberal nucleus; PAG, periaqueductal gray; PB, parabrachial nucleus; PCRt, parvicellular reticular nucleus; PHD, posterior hypothalamic area, dorsal; PLH, peduncular part of the lateral hypothalamus; Pn, pontine reticular nucleus; PPN, pedunculopontine nucleus; PVP, paraventricular thalamic nucleus, posterior; Ra, midbrain raphe; SC, superior colliculus; TH, thalamus; VMH, ventromedial hypothalamic nucleus; ZI, zona incerta. Error bars are s.e.m.; box-and-whisker plots show median, IQR, and range; shading in (M) is mean ± s.e.m.; ** *p* < 0.01.

### Glutamatergic PAG neurons project broadly to diverse brainstem and forebrain regions

VGluT2^+^ PAG neurons showed a widely distributed, ipsilaterally biased projection pattern, with the densest innervation found in the midbrain and pons, and sparser but consistent targeting of medullary and forebrain structures. Cortical and cerebellar structures were entirely spared (Figure 1F-G) (see also (21–23, 35)).

For both subdivisions of the PAG, we observed the highest density of VGluT2^+^ synaptic puncta in the PAG itself, indicating a high level of recurrent connectivity (Figure 1G). This within-PAG connectivity was strongly biased towards the injection site (and hence to the column of injection), yet it was not restricted to a single column (Figure 1H-I). Additionally, synaptic puncta from VGluT2^+^ neurons of both PAG subdivisions spread across the entire anterior-posterior extent of the PAG, significantly exceeding the vicinity of the injection site (*p* < 0.01, paired Wilcoxon signed-rank test; Figure 1J-M). Together, these data reveal extensive recurrent connectivity in the PAG, spanning across columns and anterior-posterior levels, which suggests that, in addition to projecting to downstream targets, these neurons modulate the local PAG network (see also (12, 16, 36–39)).

Outside of the PAG, we observed markedly different projection patterns for the VGluT2^+^ neurons of the dorsal and ventral divisions of the PAG. For the dPAG, the most prominent projection target of VGluT2^+^ neurons was the CnF (Figure 1F-G). In other midbrain and pontine brain regions, we observed considerably sparser synaptic densities, including the pedunculopontine, Barrington’s, laterodorsal tegmental, and pontine reticular nuclei, and locus coeruleus (Figure 1F-G). In the medulla, VGluT2^+^ dPAG neurons targeted Mc and lateral paragigantocellular nucleus (LPGi; Figure 1F-G). In the forebrain, selected lateral hypothalamic and thalamic areas were only sparsely targeted by VGluT2^+^ dPAG neurons (Figure 1F-G). These anterograde tracings from VGluT2^+^ neurons establish that the excitatory output from the dPAG is concentrated in the midbrain, with the CnF as its dominant downstream target – a projection pattern consistent with a prominent role in the midbrain defensive circuit (24, 40, 41).

In contrast to a midbrain-dominated dPAG output, VGluT2^+^ neurons of the vPAG densely innervated a larger number of brain regions, primarily in the pons and medulla (Figure 1F-G). The strongest projection density outside the PAG was found in Barrington’s nucleus (Bar) – a target only sparsely innervated by VGluT2^+^ dPAG neurons – followed by the CnF, locus coeruleus, and pedunculopontine nucleus. The most prominent medullary target of VGluT2^+^ vPAG neurons was Mc, and – to a lower extent – LPGi (Figure 1F-G). Similar to the dPAG, VGluT2^+^ vPAG neurons sparsely innervated forebrain areas, although these also included amygdalar nuclei (Figure 1F-G).

These data reveal two qualitatively distinct output patterns within the PAG: a narrow, high-density pathway from the dPAG primarily to a single midbrain locomotor structure, the CnF, versus a broader, multi-target distribution from the vPAG across pontine and medullary premotor and autonomic structures, with Bar as its most prominent target. An extended overview of PAG projection targets is shown in Figure S2.

### GABAergic PAG neurons are locally projecting with few, specific long-range targets

The projection architecture of VGAT^+^ PAG neurons differed from that of VGluT2^+^ projection neurons; they almost exclusively innervated midbrain structures, while pontine, forebrain, and medullary areas received only sparse input, consistent with a predominantly local role of GABAergic PAG neurons (16, 39, 42) (Figure 2B-C). Similar to VGluT2^+^ neurons, VGAT^+^ neurons of all columns primarily innervated the PAG itself (Figure 2C). Their synaptic distribution pattern was biased towards the column of injection (Figure 2D-E) and the synaptic spread covered the entire anterior-posterior extent of the PAG, exceeding the injection site significantly for both dPAG and vPAG (*p* < 0.01, paired Wilcoxon signed-rank test; Figure 2F-I).

**Figure 2.**
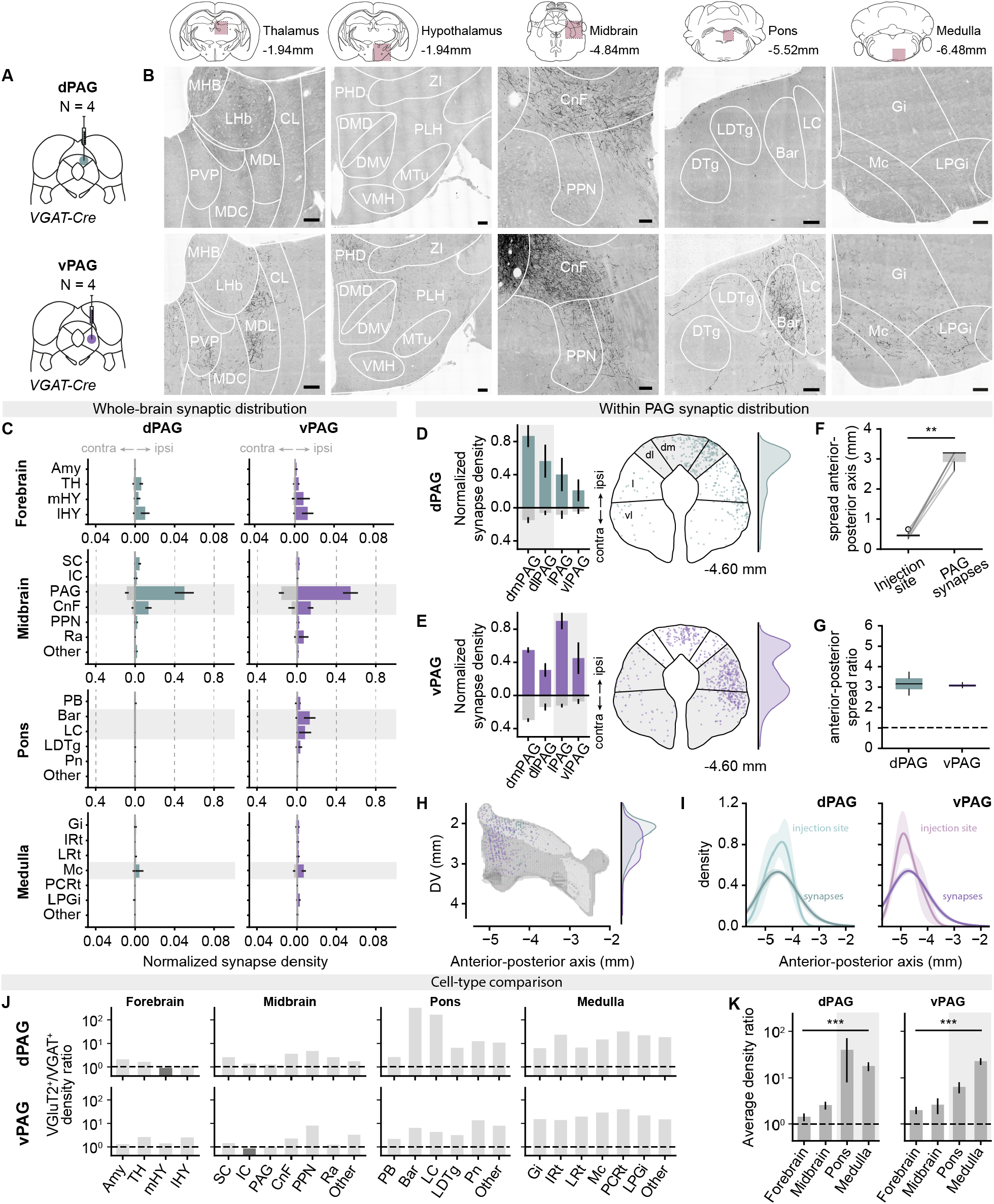
Whole-brain distribution of synaptic terminals from VGAT^+^ PAG neurons. **(A)** Cell-type- and column-specific anterograde labelling strategy for VGAT^+^ neurons from the dPAG and vPAG (top and bottom panel, respectively). **(B)** Example images of mGFP-labelled fibres in the thalamus, hypothalamus, mesencephalic locomotor region/midbrain, pons, and medulla (from left to right) of VGAT^+^ neurons from the dPAG (top panels) and vPAG (bottom panels; scale bars, 100 µm). **(C)** Regional density of synapses from VGAT^+^ dPAG (left panels, teal, *N =* 4 mice) and vPAG neurons (right panels, purple, *N =* 4 mice), normalized to the densest region per brain. Grey and coloured bars represent contralateral and ipsilateral hemispheres, respectively. The PAG and CnF are strongly innervated across columns, Bar and LC are preferentially innervated by vPAG neurons, and Mc is weakly innervated across columns (shaded areas). **(D)** Local density of synapses from VGAT^+^ neurons of the dPAG, normalized to the densest region per brain and shown for the ipsi- and contralateral hemisphere, respectively (left panel). Example spatial distribution of VGAT^+^ synapses within the PAG at one coronal level, downsampled by a factor of 10, and 2D kernel density estimation (KDE) of synapse positions (right panel). **(E)** Same as (D) but for synapses from VGAT^+^ neurons of the vPAG. **(F)** Within-PAG spread of the localization of synapses versus extent of the injection site along the anterior-posterior axis of the entire PAG, demonstrating that synaptic spread significantly exceeds the injection site (*N =* 8 mice, paired Wilcoxon signed-rank test: *p* < 0.01). **(G)** Ratio of within-PAG spread of synapses over the extent of the injection site by column of injection. The dashed line indicates a spread ratio of 1 (synaptic spread equals injection site extent). The ratio does not differ between the injection groups (Mann-Whitney U test: *p* = 0.886). **(H)** Reconstruction of synaptic distribution within the PAG from dPAG (teal) and vPAG neurons (purple), downsampled by a factor of 400, and 2D KDE of synapse positions along the dorsoventral axis. **(I)** KDE of injection site area and number of synapses along the anterior-posterior extent of the PAG by injection column (dPAG: left panel, teal; vPAG: right panel, purple). **(J)** Ratio of normalized regional density of synapses from VGluT2^+^ neurons over VGAT^+^ neurons per brain region, by column of injection (dPAG: top panels; vPAG: bottom panels). A ratio *>* 1 (above dashed line) indicates preferential targeting by VGluT2^+^ neurons (light grey); < 1 indicates preferential targeting by VGAT^+^ neurons (dark grey). **(K)** Summary of data in (J). Average density ratio across regions per major brain division, by column of injection (dPAG: left; vPAG: right; forebrain: n = 4, midbrain: n = 7, pons: n = 6, medulla: n = 7 subregions), demonstrating a higher ranking of pontine and medullary targets for VGluT2^+^ projections compared to VGAT^+^ projections from both PAG subdivisions (grey shading, Kruskal-Wallis test: *p* < 0.001 for both dPAG and vPAG, Mann-Whitney U test post-hoc comparisons). Abbreviations of brain regions are the same as in Figure 1. Error bars are s.e.m.; box-and-whisker plots show median, IQR, and range; shading in (I) is mean ± s.e.m.; ** *p* < 0.01, *** *p* < 0.001.

Outside of the PAG, VGAT^+^ neurons of both PAG subdivisions most strongly innervated the CnF (Figure 2B-C), suggesting that the CnF receives not only glutamatergic, but also direct GABAergic input from the PAG. Beyond the CnF, VGAT^+^ vPAG neurons, but not VGAT^+^ dPAG neurons, also targeted Bar (18, 43) and locus coeruleus at considerable densities, while forebrain and medullary regions were only sparsely innervated by both columns (Figure 2B-C; see Figure S2 for a full overview of projection targets).

To compare the projection patterns of VGluT2^+^ and VGAT^+^ PAG neurons, we calculated the ratio of their normalized synaptic densities for each analysed brain region (Methods). Pontine and medullary targets were more strongly represented in the VGluT2^+^ map for both the dPAG and vPAG, whereas forebrain and midbrain targets were equally represented in the two projection maps (Figure 2J-K). This suggests that glutamatergic and GABAergic PAG neurons operate on different spatial scales: glutamatergic PAG neurons both engage brainstem motor-control centres for behaviour execution and act upon the local PAG circuitry, whereas GABAergic PAG neurons are more likely to serve a primarily local, modulatory role.

### PAG neurons segregate into distinct subpopulations by projection target

The anterograde synaptic projection atlas established that glutamatergic and GABAergic PAG neurons collectively innervate a range of brain regions relevant for motor and autonomic control, but cannot resolve whether this reflects collateralizing neurons with multiple projection targets or separate subpopulations each dedicated to a single target (Figure 3A-C). To distinguish these possibilities, we performed dual-colour retrograde tracing using herpes simplex virus type 1 (HSV-1). We selected four well-characterized candidate effectors that shared a common function, received broad input from all PAG columns, or received input from the same PAG column but served distinct functions: the CnF and LPGi, both implicated in the initiation of high-speed locomotion and escape (26, 27, 44); Mc, which implements behavioural arrest including freezing (16, 27); and Bar, which controls micturition and, like Mc, receives input predominantly from the vlPAG (43) (Methods; Figure S3 and S4A-C).

**Figure 3.**
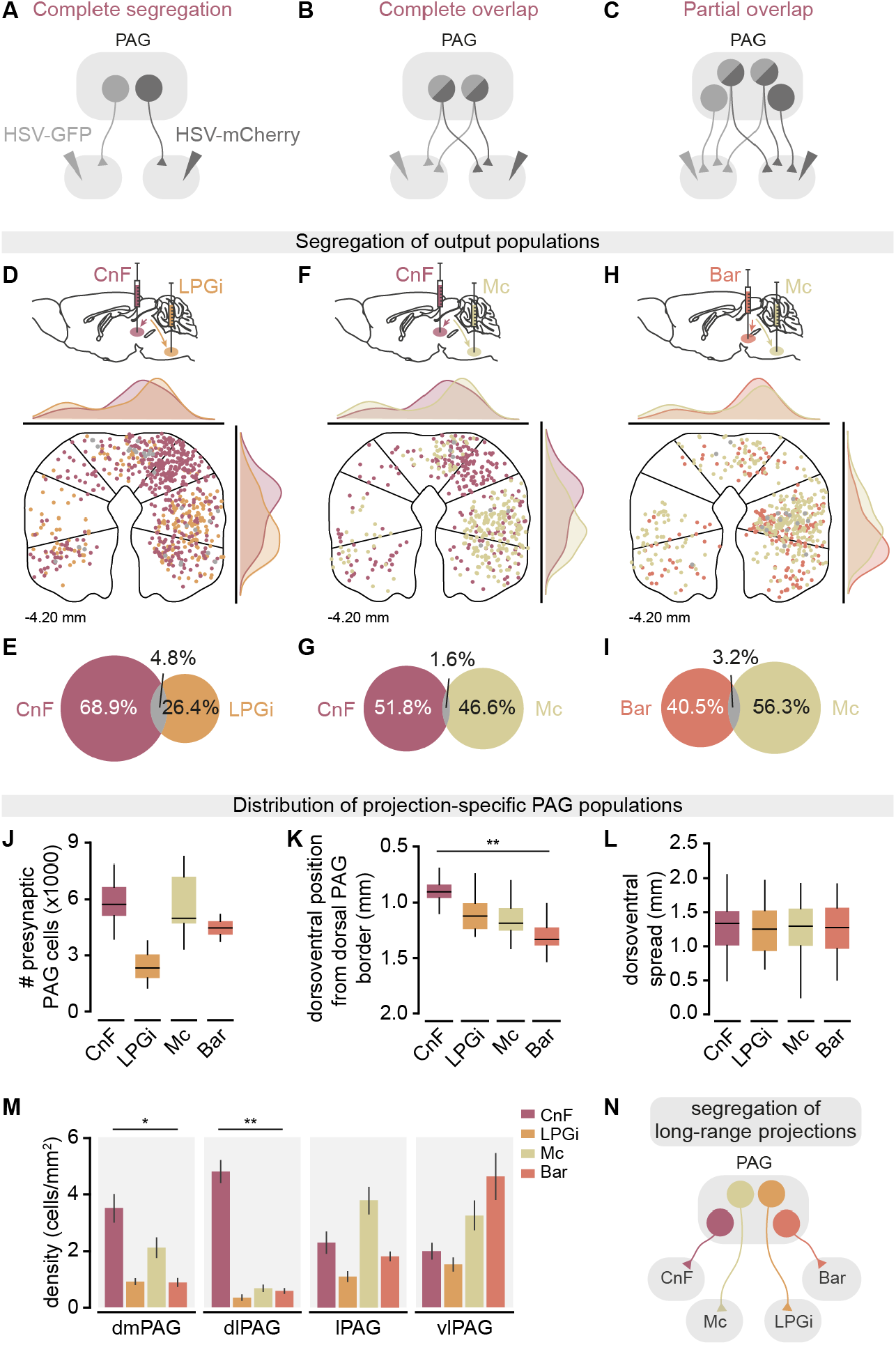
Neurons in the periaqueductal gray segregate by projection target. **(A-C)** Schematics of hypothetical population architectures: **(A)** complete segregation, in which distinct PAG neurons project exclusively to each target; **(B)** complete overlap, in which all PAG neurons project to both targets; **(C)** partial overlap, in which subpopulations project to one or both targets. **(D)** Dual-colour retrograde labelling strategy from CnF and LPGi (top panel). Example distributions of retrogradely labelled cell bodies in the PAG at one coronal level, downsampled by a factor of 5, and 2D KDE of the distribution of PAG neurons per projection target along the mediolateral and dorsoventral axis of the PAG (bottom panel). **(E)** Percentages of retrogradely labelled PAG cells expressing the fluorescent markers for CnF- and LPGi-projecting neurons (*n* = 63 slices in *N =* 3 mice). **(F)** Same as (D) from CnF and Mc. **(G)** Same as (E) for CnF- and Mc-projecting neurons (*n* = 43 slices in *N =* 3 mice). **(H)** Same as (D) from Bar and Mc. **(I)** Same as (E) for Bar- and Mc-projecting neurons (*n* = 37 slices in *N =* 2 mice). **(J)** Total number of presynaptic PAG neurons by projection target, estimated from stereological counts in sampled sections, corrected for section sampling interval, which did not differ significantly between projection targets (Kruskal-Wallis test: *p* = 0.085). **(K)** Average dorsoventral coordinates relative to the dorsal PAG border of retrogradely labelled PAG neurons with significant differences between projection targets (linear mixed model (Wald F): *p* < 0.01, t-test post-hoc comparisons). **(L)** Spread of PAG neurons in the dorsoventral axis by projection target. No significant difference was observed between the groups (linear mixed model (Wald F): *p* = 0.225). **(M)** Density of retrogradely labelled neurons per PAG column by projection target. CnF-projecting neurons were significantly enriched in the dmPAG and dlPAG relative to the other projection populations (*n =* 126 slices in *N =* 8 mice, linear mixed model (Wald F): *p* < 0.05 (dmPAG), *p* < 0.01 (dlPAG), *p* = 0.164 (lPAG), *p* = 0.131 (vlPAG); t-test post-hoc comparisons). **(N)** Schematic summary of the observed ‘labelled line’ projection logic, in which distinct PAG subpopulations project selectively to individual downstream targets. Bar, Barrington’s nucleus; CnF, cuneiform nucleus; LPGi, lateral paragigantocellular nucleus; Mc, magnocellular reticular nucleus; PAG, periaqueductal gray. Across the figure, the following colours designate the respective projection targets: CnF, bordeaux; LPGi, orange; Mc, ochre; Bar, salmon. Error bars are s.e.m.; box-and-whisker plots show median, IQR, and range; * *p* < 0.05, ** *p* < 0.01.

Dual tracing from CnF and LPGi revealed that the majority of labelled PAG neurons (68.9 ± 4.1%) projected exclusively to the CnF (PAG_CnF_), while 26.4 ± 3.8% projected exclusively to LPGi (PAG_LPGi_), and only 4.8 ± 1.4% of neurons projected to both CnF and LPGi (Figure 3D-E and S4D). Second, we compared PAG_CnF_ and Mc-projecting neurons (PAG_Mc_), two populations whose downstream targets initiate mutually exclusive defensive behaviours – escape and freezing, respectively (16, 24). While we observed a similar percentage of PAG_CnF_ and PAG_Mc_ neurons (51.8 ± 5.6% and 46.6 ± 5.9%, respectively), the fraction of neurons projecting to both regions simultaneously was only 1.6 ± 0.3% (Figure 3F-G and S4E). Third, we compared PAG_Mc_ neurons to neurons projecting to Bar (PAG_Bar_), two regions predominantly targeted by vlPAG neurons (Figure 1G and 2C). Here, of all labelled PAG cell bodies, 56.3 ± 10.3% projected exclusively to Mc, 40.5 ± 10.2% to Bar, and only 3.2 ± 0.2% to both regions (Figure 3H-I and S4F). Across all pairings, the observed overlap was significantly smaller than expected if projections to the two targets were assigned independently, and remained so after correcting for the technical ceiling of viral co-expression within one region (*p* < 0.05 for both comparisons, pairwise Wilcoxon signed-rank test; Methods; Figure S4G-J). These results support a model in which PAG long-range projections are organized into largely segregated, target-specific subpopulations rather than collateralizing onto multiple downstream regions.

We next assessed the spatial distribution of the target-specific PAG subpopulations. The total number of retrogradely labelled neurons in the PAG was comparable across all four target regions (*p =* 0.085, Kruskal-Wallis test; Figure 3J and S4K), and we observed a similar distribution of labelled neurons across the anterior-posterior and mediolateral axes of the PAG, with the majority of neurons found in the caudal PAG, ipsilateral to the injection site (Figure S4L-M). However, we found a significant difference in the columnar distribution of PAG neurons based on their projection target, in line with our anterograde tracing results. The majority of PAG_CnF_ neurons were located in the dorsal and lateral parts of the PAG, while the majority of PAG_Bar_, PAG_LPGi_, and PAG_Mc_ neurons were located more ventrally (*p* < 0.01, linear mixed model (Wald F); Figure 3K). Yet, none of these populations were confined to a single column; each spanned the full dorsoventral extent of the PAG (Figure 3L). Notably, while PAG neurons projecting to each of the four brain regions were distributed across columns with significant dorsoventral gradients, neurons in the dorsolateral column of the PAG projected almost exclusively to the CnF (*p* < 0.01, linear mixed model (Wald F); Figure 3M).

To determine the percentage of GABAergic PAG neurons among the retrogradely labelled populations, we immunohistochemically stained against GABA (Figure S4N). GABA^+^ neurons contributed to all four projection populations, with the PAG_Mc_ population containing a significantly smaller percentage of GABA^+^ neurons (*p* < 0.05, linear mixed model (Wald F); Figure S4O). The density distribution of GABA^+^ neurons across PAG columns revealed the same pattern observed for the entirety of the retrogradely labelled neurons: GABA^+^ PAG_CnF_ neurons were mainly observed in the dmPAG, dlPAG, and lPAG, while GABA^+^ PAG_Bar_, PAG_LPGi_, and PAG_Mc_ neurons localized more prominently in the lPAG and vlPAG (Figure S4P). These findings suggest that the anatomical segregation we observe is not attributable to differential cell type composition – both glutamatergic and GABAergic neurons contribute to each target-specific output population. Together, these findings support a ‘labelled line’-like organization of PAG output, in which non-overlapping subpopulations project selectively to individual downstream targets (Figure 3N), and in which the CnF-specialized dlPAG represents the clearest example of column-level target specificity.

### CnF-projecting PAG neurons form local collaterals within the PAG

The anterograde tracing revealed dense local connectivity within the PAG (Figure 1M and 2I), but could not answer whether the described output populations contribute to this. To address this, we focused on the dPAG projections to the CnF (dPAG_CnF_) as the CnF is the main output target of the PAG and the dPAG_CnF_ pathway is functionally defined: dPAG glutamatergic neurons control escape initiation and vigour (12, 39), and projection-specific activation of dPAG_CnF_ neurons is sufficient to drive flight (24).

First, we analysed the topography of dPAG synapses in the CnF to test whether this projection has any observable gradients in synapse distribution. The distribution of synaptophysin-labelled puncta in the CnF from VGluT2^+^ and VGAT^+^ dPAG neurons was uniform and did not differ along the mediolateral and anterior-posterior axes of the CnF (Figure 4A-B), with significantly higher synapse counts in VGluT2-Cre than in VGAT-Cre mice (*p* < 0.05, Mann-Whitney U test; Figure S5A-B). Second, to test these connections functionally, we unilaterally injected HSV-Cre into the CnF and AAV-DIO-ChR2 into the dPAG of wildtype mice, to express ChR2 exclusively in dPAG_CnF_ neurons, and performed patch-clamp recordings from identified, ChR2-negative neurons in the dPAG and CnF in acute midbrain slices *ex vivo* (Figure 4C-D and Figure S5C). Using full-field illumination with blue light (470 nm) to test for the presence of light-evoked excitatory postsynaptic currents (eEPSCs), we found that dPAG_CnF_ neurons project locally within the dPAG with a high connection probability (77.8%) and a slightly lower connection probability to the CnF (66%; Figure 4E). In addition to similar amplitudes, median synaptic latencies of dPAG_CnF_-CnF and dPAG_CnF_-PAG eEPSCs of ∼1.60 ms suggest a predominantly monosynaptic connection to each target (Figure 4F). The paired pulse ratio (PPR) across a five-pulse train, delivered at 20 Hz and compared to the first pulse, suggests that both connections allow for the integration of synaptic inputs over time, with a facilitating PPR for dPAG_CnF_-CnF (Figure 4G). Accordingly, in five out of nine CnF cells that fired action potentials in response to light stimulation, the spiking probability was similar across the pulse train (Figure 4H). Next, we tested the percentage of neurons that received evoked inhibitory postsynaptic currents (eIPSCs) and found that 44% of dPAG neurons and 43% of CnF neurons received inhibitory input (Figure 4I). The amplitude of dPAG_CnF_-CnF eIPSCs was significantly smaller than that of dPAG_CnF_-PAG eIPSCs (*p* < 0.05, Mann-Whitney U test) and we observed a skewed distribution of the synaptic latency for dPAG_CnF_-CnF eIPSCs in comparison to dPAG_CnF_-PAG eIPSCs, with latencies of up to 9.2 ms (*p* < 0.05; Mann-Whitney U test; Figure 4J). The relative latencies between eEPSCs and eIPSCs within individual CnF neurons showed a median latency of 0.8 ms with a range of 8.3 ms (Figure 4K). While these data support the presence of direct inhibitory dPAG_CnF_-CnF synaptic connections, they suggest that both direct and multi-synaptic feed-forward inhibition shape information transmission in the CnF. In contrast, the same dPAG_CnF_ neurons provide more balanced excitatory-inhibitory transmission within the dPAG with similar eIPSC and eEPSC amplitudes and latencies (*p* = 0.451 and *p* = 0.845, Mann-Whitney U test). Together, these data support a dual role for dPAG_CnF_ neurons in shaping downstream locomotor output in a target-specific manner, while at the same time shaping ongoing PAG activity through recurrent excitatory and inhibitory synaptic motifs (Figure 4L).

**Figure 4.**
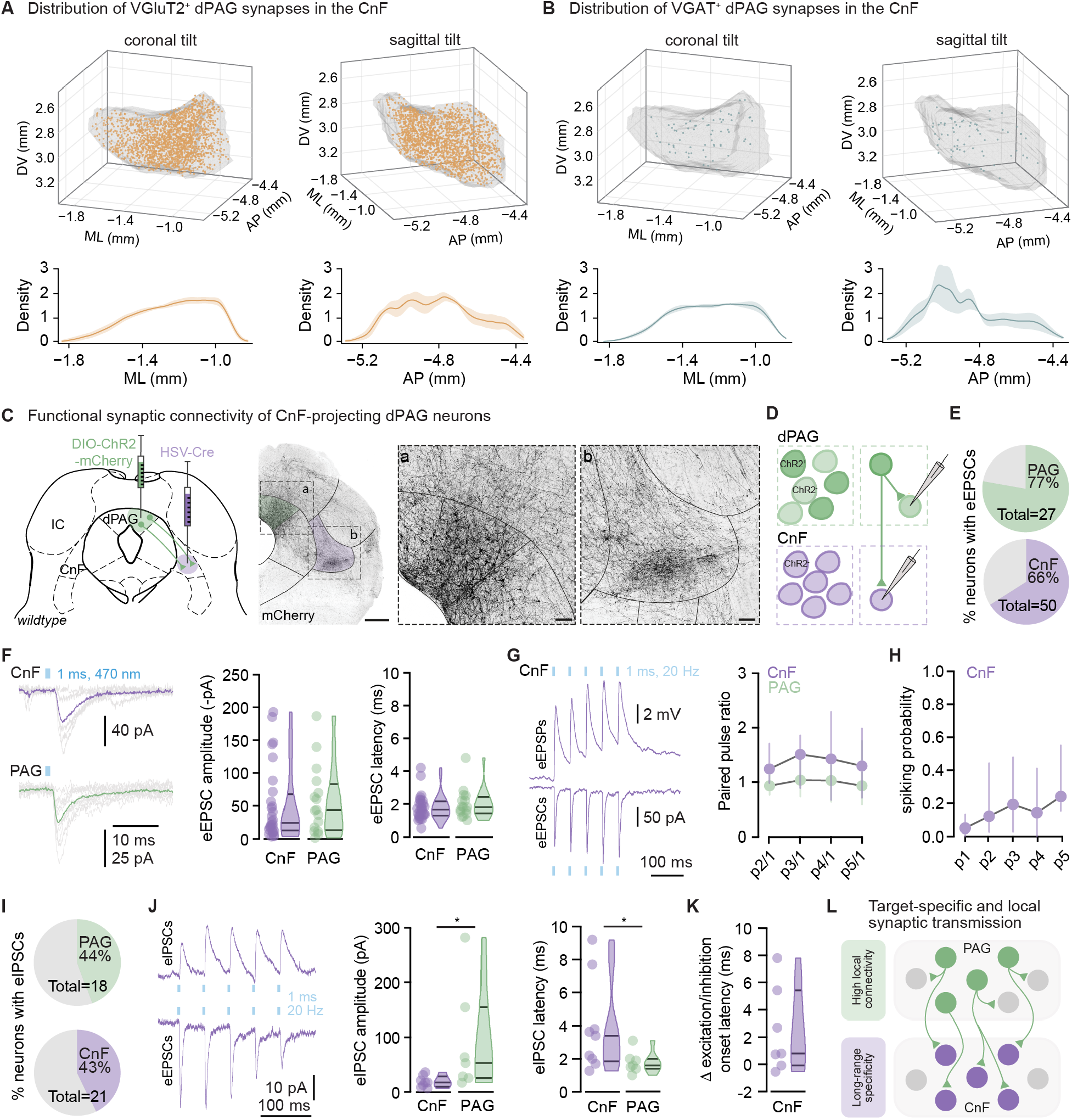
CnF-projecting dPAG neurons form local collaterals within the dPAG. **(A)** Distribution of VGluT2^+^ dPAG synapses in the CnF along the mediolateral (coronal tilt) and anterior-posterior (sagittal tilt) axes (top panels; *N =* 4 mice, 48747 ± 16194 synapses), downsampled by a factor of 100, and 2D KDE of synaptic density along each axis (bottom panels). Density showed no spatial gradient along either axis (*p* = 0.587 for both, one-sample t-test). **(B)** Same as (A) for VGAT^+^ dPAG synapses (*N =* 4 mice, 2079 ± 1186 synapses; *p* = 0.658 for both, one-sample t-test). **(C)** Experimental design for ChR2 expression restricted to dPAG_CnF_ neurons (left panel) and example histology of ChR2-mCherry expression (dPAG, green; CnF, purple), showing somata and fibres in the dPAG (a) and fibres in the CnF (b) (middle and right panel; scale bars: 500 µm, left, 100 µm, middle and right). **(D)** ChR2 expression pattern and patch-clamp recording configuration in acute coronal slices. Both local connections within the dPAG and connections to the CnF are tested using CRACM. **(E)** Connection probability for dPAG_CnF_-PAG (*n* = 21 of 27 neurons, top panel) and dPAG_CnF_-CnF (*n* = 33 of 50 neurons, bottom panel), tested in voltage clamp for eEPSCs. Of *N* = 8 animals tested in total, data for both CnF and dPAG were acquired in four animals, and two animals each have data for only CnF or dPAG; connectivity rates did not differ and were pooled (CnF: *p* = 0.534, dPAG: *p* = 0.662; Fisher’s exact test). **(F)** Individual (grey) and average eEPSC for an example CnF (top) and dPAG neuron (bottom; left panel). Median eEPSC amplitude (middle panel) and synaptic latency (right panel) did not differ between CnF and dPAG neurons (*n* = 28 and 16 cells, *N* = 6 mice each, four shared between both groups; *p* = 0.552 and *p* = 0.373, Mann-Whitney U test). **(G)** Average eEPSC and light-evoked excitatory postsynaptic potential (eEPSP) of an example CnF neuron during a pulse train (5 pulses, 1 ms at 20 Hz; left panel) and median paired pulse ratio of each peak relative to the first pulse across the pulse train for CnF (*n* = 24 cells, purple) and dPAG (*n* = 14 cells, green; *N* as in (F); right panel). **(H)** Median spiking probability of a subset of CnF neurons recorded in current clamp shown across the pulse train for each pulse (p). **(I)** Same as (E) for eIPSCs: *n* = 8 of 18 cells for dPAG_CnF_-PAG (green, top panel) and *n* = 9 of 21 cells for dPAG_CnF_-CnF (purple, bottom panel). **(J)** Average eIPSC (top) and eEPSC (bottom) of an example CnF neuron recorded at 0 mV and -70 mV, respectively (left panel). Median eIPSC amplitude and synaptic latency for connected CnF (*n* = 8 and 9 cells; one cell excluded from amplitude analysis as it contained an IPSC-EPSC sequence) and dPAG neurons (*n* = 7 cells, right panels; *p* = 0.029 and *p* = 0.034, respectively; Mann-Whitney U test). **(K)** Within-cell difference in the relative latency between eIPSCs and eEPSCs for CnF neurons (*n* = 7 cells, *N* = 4 mice). **(L)** Schematic summary of collateralization of CnF-projecting dPAG neurons. Shading in (A, B) is mean ± s.e.m.; violin plots show median, IQR, and range; point-range plots show median and IQR; p1-p5, pulses; * *p* < 0.05.

## Discussion

We establish a whole-brain projection atlas of the PAG at column-, synapse- and cell-type-resolution, revealing a differential dorsal and ventral output organization with predominantly glutamatergic long-range projections and dense recurrent glutamatergic and GABAergic connectivity. Retrograde tracing from four motor-related targets revealed that PAG neurons are organized into non-overlapping, target-specific output populations – a ‘labelled line’ architecture in which most neurons project selectively to single downstream effectors rather than broadcasting to multiple targets. Using CRACM, we further show that CnF-projecting PAG neurons branch locally and project to other PAG neurons at a high rate. Together, our findings provide a circuit basis for how the PAG generates distinct behavioural outputs from spatially intermingled sets of neurons and support a model in which distinct output populations operate at two spatial scales: long-range projections to dedicated, action-specific targets, and local connections that modulate the within-PAG circuitry.

### A segregated output architecture for PAG-mediated behaviours

Our data reveal an organizational principle in which PAG neurons projecting to different motor targets form segregated, target-specific output populations, and quantify it across four motor-related targets – CnF, LPGi, Mc, and Bar (see also (33)). Co-labelling rates across all pairings indicate near-complete population segregation. This degree of target specificity, observed consistently across pairings with similar (CnF and LPGi) and opposing functions (CnF and Mc), as well as shared anatomical columnar input but different functions (Mc and Bar), supports a generalized ‘labelled line’ principle rather than a pairing-dependent feature. The functional significance of this architecture is well illustrated in the vlPAG. Optogenetic stimulation of glutamatergic vlPAG neurons can elicit both freezing and micturition, whereas stimulation of glutamatergic vlPAG terminals in Mc and Bar elicits only freezing and micturition, respectively (16, 18). Our data provide the anatomical basis for this observation: Mc- and Bar-projecting PAG populations are of approximately equal size and, while they overlap spatially, they are almost entirely non-overlapping at the single-neuron level. This suggests that spatially intermingled neurons within a single column drive functionally distinct behavioural outputs through dedicated projections to a single output region rather than axonal collaterals simultaneously targeting multiple effectors. Whether this projection motif holds across all domains, both locomotor and autonomic, and whether local recurrent connectivity within the PAG allows different pathways to synchronize their output to coordinate multifaceted behavioural responses, such as integrating cardio-behavioural responses during defensive freezing (15), remains an important open question.

While the columnar organization of the PAG remains valid as a broader description of PAG functional topography, we show that target identity provides an important additional organizational axis that is partially correlated with, but not reduced to, columnar identity. Single-cell transcriptomic profiling of the PAG identified over 140 molecularly distinct neuronal populations forming spatial metaclusters that align with, but are not limited to, the classical column boundaries (45), and genetically defined subpopulations can oppose the behaviour predicted by their columnar identity (46). The PAG’s functional architecture is thus better understood as the superposition of several partially independent organizational axes – columnar, molecular, and, as we show here, target-defined.

### Modular output as a conserved principle of motor circuits

An organizational logic with target-defined output populations has been reported in motor cortex, basal ganglia, midbrain, and spinal cord, where intermingled, functionally dedicated populations project to distinct targets (25, 27–29, 31, 47). Our findings place the PAG within this group and suggest that a modular output organization is a conserved feature of motor-related circuits. Such an architecture offers two main advantages: first, output channels can be recruited independently, so that one output is engaged without driving another (25, 27). Whether each of these populations is innervated by a distinct set of afferents will have to be established by multi-synaptic, projection-specific retrograde tracing. Second, segregated output populations can be modulated individually (48), allowing internal state, prior experience, or neuromodulatory tone to bias which behaviour is selected – a plausible mechanism for the state-dependence of instinctive behaviours (49). While the main behavioural output of some of these target-defined populations has been established using projection-specific stimulation studies (16, 18, 24, 26, 27, 44), our anatomical data cannot resolve the level at which they act. A pathway could command a complete behavioural program, such as a shelter-directed escape, or it could encode a single action or variable that is recruited across behavioural contexts (e.g., *speed* through LPGi projections). In the latter case, the target could specify which action is executed and the column (and molecular identity of the neurons) could provide the ‘behaviour context’ in which it is executed. For example, CnF-projecting neurons might initiate fast locomotion and could be recruited from molecularly defined neurons predominantly located in the dPAG during escape and in the lPAG during predatory pursuit. Recordings from projection-defined PAG populations and their downstream targets across behavioural contexts in the *same* animal should distinguish these possibilities (15, 50, 51).

### Local versus long-range organization of the periaqueductal gray

In addition to a target-specific organization, we provide evidence for cell-type-specific projection patterns. VGluT2^+^ neurons broadly projected across the forebrain and brainstem, with dense innervation of midbrain, pontine, and medullary targets. VGAT^+^ neurons, however, were primarily locally projecting, with only sparse long-range outputs. These observations align with the demonstrated role of tonically active GABAergic dPAG neurons in locally inhibiting excitatory PAG output (39, 42), and in general, with a disinhibitory motif for output control in the PAG (16, 39), suggesting local inhibition as the primary output mode of GABAergic PAG neurons. At the same time, long-range PAG output neurons are not exclusively long-range: we show that CnF-projecting PAG neurons strongly collateralize within the PAG with a high connection probability, allowing them to modulate the PAG-internal circuitry while simultaneously driving their downstream target. This positions PAG_CnF_ neurons as both output and local circuit elements (see also (52)). Whether every neuron in a target-defined population collateralizes locally, or whether projection-only neurons exist alongside mixed neurons, remains open.

At the synaptic level, PAG_CnF_ neurons engage the downstream target and the local PAG circuit through distinct synaptic motifs. While short-latency excitatory synaptic transmission was evoked in the majority of PAG and CnF neurons with similar amplitudes, inhibitory currents were recorded in less than half of PAG and CnF neurons and showed region-specific latency distributions and amplitudes. Within the PAG, inhibitory synaptic latencies were short, while latencies in the CnF were more distributed and extended well beyond the monosynaptic range, possibly indicating a recruitment of CnF GABAergic neurons. A single output channel can therefore deliver feed-forward inhibition downstream while also providing direct local inhibition – two motifs operating on different timescales, so that the local circuit is shaped before the downstream computation is resolved. The PAG_CnF_ population also showed a differential excitation-inhibition balance in the two targets: in the CnF, inhibitory currents were substantially smaller than excitatory ones, whereas in the PAG the two matched in amplitude and latency. This suggests that at the downstream target excitation dominates, as expected from an output channel built to drive a behaviour, while excitation and inhibition are balanced to regulate the local PAG circuitry. PAG_CnF_-CnF synapses facilitated across a 20 Hz train, consistent with a pathway built to sustain an action. Importantly, neurons in the dPAG increase their firing rate by ∼17 Hz at escape onset and sustain this elevated firing for the duration of the escape, with the duration of the depolarization predicting the duration of the behaviour (53). A facilitating synapse driven at close to this rate (54), with little opposing inhibition, would transmit such a maintained drive reliably – converting a sustained PAG command into sustained CnF recruitment. Notably, the CnF and PAG both receive input from the superior colliculus and the CnF also projects back to the PAG (26, 40, 55–58). Thus, an important next step will be to resolve how these recurrent loops interact with the motifs described here and what their functional role is.

Local collateralization further provides an essential mechanism of mutual control between the modular output populations of the PAG: the activity of one output population is conveyed to the local PAG network at short latency. The functional significance of this depends on the cellular identity, and projection target, of the postsynaptic partner. If, for example, the collaterals recruit local GABAergic neurons, these could inhibit other output populations, so that the recruitment of one channel would suppress competing channels and enforce mutual exclusivity of the behaviours they drive. Whether this constitutes a mechanism for behaviour selection remains an important open question. If instead neurons of the same channel are excited, this could provide recurrent amplification and thereby a mechanism for persistent activity to support a selected behaviour on the scale of seconds (12). Given the recruitment-based scaling described above, within-pathway collaterals could scale behaviour by activating additional neurons rather than by strengthening individual responses (53). These possibilities are not mutually exclusive and could operate in parallel. In addition, behaviours with concurrent motor and autonomic responses (e.g., locomotor arrest, increased muscle tone and bradycardia during freezing) could be ‘assembled’ through the co-recruitment of appropriate PAG populations through feed-forward excitatory connections. To assess this, a detailed anatomical microcircuit analysis will help to answer whether the PAG contains dedicated subnetworks of neurons that process incoming sensory and internal inputs, drive motor and autonomic commands, shape ongoing activity through recurrent excitatory and inhibitory synaptic motifs, and allow for the selection of competing instinctive behaviours through lateral inhibition motifs (12, 16, 59).

Together, our data show that the PAG contains a modular output architecture to support a range of complex instinctive behaviours. Long-range output is organized into segregated, target-defined output populations that are spatially intermingled across, and partially independent of, the columnar structure. This organization explains how spatially overlapping neurons can drive distinct and mutually exclusive behaviours. These output populations are further embedded in a dense local circuitry, through which the activity of one population can influence others. We propose that this modular organization with local modulation provides a general solution by which premotor circuits select, scale, and combine behavioural output. The atlas and projection-defined populations described here provide the anatomical entry points to test this directly. Recording and manipulating identified output populations across behavioural contexts should reveal how the PAG translates sensory and internal state information into the selection and execution of instinctive behaviours, and their associated autonomic and endocrine components.

## Supporting information

Supplemental_Statistical_Reporting

## Acknowledgements

This work was funded by the Max Planck Society, the Behrens-Weise Foundation, and by a European Research Council grant (project 101164752). We thank the animal facility at the Max Planck Institute for Brain Research for animal care and G. Rosiles and R. Pedro da Silva for technical assistance. We thank the members of the Stempel lab for discussions and P. Tovote, S. Keshavarzi, D. Evans, P. Brand and G. Laurent for comments on the manuscript. This manuscript was typeset using a modified version of the HenriquesLab bioRxiv template.

## Author Contributions

A.S. and A.V.S. conceived the project and designed experiments. A.S. performed viral tracings. A.T. and A.V.S. performed *ex vivo* electrophysiology. J.N.M. wrote and applied code for injection site determination. A.S. and E.S.D.B. conducted immunohistochemical stainings and tracing quantification. A.S., A.T., and A.V.S. analysed data. A.V.S. supervised the project. A.S. and A.V.S. wrote the manuscript, and all authors read and commented on the manuscript.

## Methods

### Animals

Adult (6–14 weeks old) male and female VGluT2-Cre (Jackson Laboratory, JAX number: 016963), VGAT-Cre (Jackson Laboratory, JAX number: 016962), and C57BL/6J wildtype (Jackson Laboratory, JAX number: 000664) mice were housed in individually ventilated cages on a 12 h light/dark cycle with *ad libitum* access to water and food. VGAT-Cre::R26-EYFP mice were generated by crossing VGAT-Cre (Jackson Laboratory, JAX number: 016962) with the Cre-dependent reporter line R26-EYFP (Jackson Laboratory, JAX number: 006148). All animal experiments were conducted in accordance with the European Union Directive 2010/63/EU on the protection of animals used for scientific purposes, the German Animal Welfare Act (Tierschutzgesetz, TierSchG), and approved by the responsible local authority (Regierungspräsidium Darmstadt; animal licence F126/2008).

### Viruses

The following viruses were used in this study and are referred to by abbreviations in the text. For anterograde tracing experiments, AAV-hSyn-FLEX-mGFP-2A-Synaptophysin-mRuby was obtained from Addgene (plasmid #71760) and packaged into an AAV2/2 serotype (#BA-109e, 1.21×10^12^vg/ml) by Charité Vector Core Facility (VCF). For retrograde tracing experiments, HSV-hEF1α-mCherry-IRES-flpo (2×10^9^vg/ml) and HSV-hEF1α-EGFP (2×10^9^vg/ml) were acquired from NeuroTools Viral Vector Core, University of North Carolina (NeuroTools, UNC). AAV2/2-CAG-eBFP2.NLS-minWPRE, acquired from VCF (#BA-376a, 5.58×10^12^vg/ml), was used as an injection site marker. For *ex vivo* optogenetic activation, HSV-hEF1α-Cre (2×10^12^vg/ml, NeuroTools, UNC) and AAV2/2-EF1α-DIO-hChR2(H134R)-mCherry (4×10^12^vg/ml, UNC Vector Core) were used.

### Surgical procedure

Drugs were prepared on the day of surgery, diluted in sterile 0.9% NaCl saline and warmed before use. Buprenorphine (0.05 mg/kg) and Metacam (5 mg/kg) were administered subcutaneously 30 min before the surgical procedure for perioperative analgesic care. Animals were anaesthetized with isoflurane in oxygen (4% initially, then 1–2% for maintenance, 1 L/min). The hair on the scalp was shaved using a trimmer (Moser) and the skin disinfected. The skin was then cut to expose the skull and Mepivacaine (5 mg/kg) was applied for additional local analgesia. A large craniotomy was made using a hand-held dental drill with a 0.3, 0.5, or 0.7 mm stainless steel burr (Fine Science Tools) to expose the area of the transverse and sagittal sinus and the anatomical border between the superior and inferior colliculi (AP from lambda: +0.1 to -1.0 mm; ML: ±1.0 mm). A durotomy was performed using a 30G hypodermic needle. The connective tissue covering the site of injection was gently pulled anteriorly using fine forceps to reveal the underlying superior colliculi. Viral vectors were delivered at ∼20 nl/min using a pulled glass pipette (10 µl Wiretrol II with a Kopf Instruments Needle/Pipette Puller, Model 720) mounted on a stereotaxic injector (Janelia Research Campus, Ronal Tool Company) connected to an oil hydraulic micromanipulator (MO-10, Narishige) on a stereotaxic frame (Model 942, Kopf Instruments). The needle was maintained at the site of injection for 10 min post injection before being retracted. The skin was sutured (6-0, Vicryl Rapide). Coordinate measurements are given from the anatomical border between superior and inferior colliculi (in AP and ML) and the skull surface (in DV).

### Anatomical tracing

#### Viral injections

For anterograde tracing, AAV-hSyn-FLEX-mGFP-2A-Synaptophysin-mRuby (50 nl) was unilaterally injected into the dPAG (anterior-posterior, AP: 0.4–0.5; mediolateral, ML: 0.1–0.5; dorsoventral, DV: 2.0–2.1) or vPAG (AP: 0.4–0.5; ML: 0.5–0.6; DV: 2.5–2.7). For dual retrograde tracing, HSV-hEF1α-mCherry-IRES-flpo and HSV-hEF1α-EGFP (100 nl each) were respectively injected at one of the target sites: CnF (AP: 0.3; ML: 1.27; DV: 3.0), Bar (AP: -0.3; ML: 0.75; DV: 3.7), Mc (AP: -1.0; ML: 0.25; DV: 5.9), LPGi (AP: -1.0; ML: 0.75; DV: 6.1).

#### Perfusion

Three weeks after viral injection, mice were anaesthetized with pentobarbital (400–800 mg/kg, final volume: 0.1 ml per 10 g, diluted 1:1 in sterile 0.9% NaCl solution), transcardially perfused with phosphate-buffered saline (PBS), followed by 4% paraformaldehyde (PFA) in PBS. Brains were dissected, post-fixed overnight in 4% PFA at 4 °C, and incubated in 30% sucrose in PBS for at least 24 h before cryopreservation. Brains were embedded in Cryo-Embedding Compound (PELCO, Ted Pella, Inc.), frozen on dry-ice, and sliced on a cryostat at 30 µm thickness (coronal sections). Floating sections were collected in cryoprotectant solution (25% glycerol, 25% ethylene glycol, 50% PBS) and stored at -20 °C. Immunohistochemically stained slices were imaged using an Axioscan Z1 light microscope (Zeiss). Example images for visualization were imaged using an LSM 880 Examiner (Zeiss).

#### Anatomical reconstruction

Images from 30 µm thick sections encompassing the entire brain along its anterior-posterior axis were acquired using a 20x objective on an Axioscan Z1 light microscope (Zeiss). Images were aligned to the Allen CCFv3 using Aligning Big Brains & Atlases (ABBA) (60) in ImageJ. Virally expressed markers were detected automatically using subcellular segmentation or manual labelling in QuPath (version 0.4.4). Automatic detections were visually validated. Injection site boundaries were delineated by applying Otsu’s automatic thresholding method to fluorescence images, which determines an optimal intensity threshold by minimizing intra-class variance of the pixel intensity histogram (61). The resulting binary masks were used to extract depth profiles along the dorsoventral axis and visually curated. Mice were clustered using hierarchical agglomerative clustering with Ward linkage and Euclidean distance, applied to KDE-smoothed depth profiles of the injection sites. The dendrogram was cut to yield a fixed number of clusters (*k* = 3), as no automated stopping criterion was applied. The same hierarchical clustering was applied to the synaptic density profiles and the optimal number of clusters (k) was determined using the silhouette score and elbow plot. The stability of cluster assignment was evaluated by bootstrap resampling (Adjusted Rand Index for k = 2, VGluT2-Cre: 0.80 ± 0.31, VGAT-Cre: 0.52 ± 0.47). All clustering and validation analyses were implemented in Python using SciPy and scikit-learn.

#### Quantification

The number and coordinates of detected objects were extracted and used for further analysis and plotting using custom-written code in Python 3.13. For the anterograde tracings, synaptic density was calculated as the number of synaptophysin-labelled puncta within a brain region divided by that region’s counted area from the Allen CCFv3, and normalized per animal to the region with the highest density, yielding values between 0 and 1. For each analysed brain region, we calculated the ratio of the normalized synaptic density of VGluT2^+^ over VGAT^+^ neurons. Because normalized synaptic densities reflect the relative projection density within a brain, this ratio measures the relative weight of a region within each projection map rather than a difference in absolute synapse numbers (ratio > 1: over-represented in the VGluT2^+^ map; < 1: over-represented in the VGAT^+^ map; = 1: equal representation). Tests for spatial gradients within the CnF were conducted by calculating the enrichment of synaptic puncta counts per mouse relative to the CnF volume in the Allen CCFv3 in 100 µm bins along each axis and fitting a per-mouse linear slope, which was tested against zero using one-sample t-tests with Benjamini-Hochberg correction across axes. For retrograde tracings, the total size of each retrogradely labelled population was estimated by multiplying the counted cell number per animal with the section sampling interval. Cell density was calculated as the number of retrogradely labelled somata within the PAG divided by the area from the Allen CCFv3. For each pairing, the overlap fraction expected under independent assignment of projection targets was calculated as the product of the two single-labelled cell fractions, and compared with the observed overlap fraction per animal. To test for the technical ceiling of viral co-expression, both vectors were co-injected into the same target region, for each tested target region, yielding an overall co-expression efficiency of 61.0 ± 3.5% (Figure S4G-I). Control stainings using the GABA antibody in VGAT-YFP tissue yielded a proportion of YFP^+^/GABA^+^ double-labelled neurons of 24 ± 3%, with a low false-positive rate of 4 ± 1% of all GABA^+^ neurons (*n* = 6 slices in *N* = 2 mice), demonstrating an expected underestimation of GABAergic neurons using GABA antibodies.

### Electrophysiological recordings in acute brain slices

#### Surgery

Dual HSV-Cre and AAV-DIO-ChR2 injections were made in the CnF and PAG of male and female C57BL/6 mice (aged 6–8 weeks). Specifically, HSV-hEF1α-Cre was injected into the CnF unilaterally (AP: 0.3; ML: 1.27; DV: 3.0; 200 nl) and AAV2/2-EF1α-DIO-hChR2(H134R)-mCherry was injected into the dPAG of the same hemisphere (AP: 0.35; ML: 0.2; DV: 2.0; 200–250 nl). This approach restricts ChR2 expression to Cre-positive PAG neurons that project to the CnF. Acute coronal slices were prepared 2–4 weeks after viral injection, with the animals aged 8–12 weeks at the time of recording.

#### Data acquisition

Mice were deeply anaesthetized with isoflurane and decapitated. Brains were quickly removed and transferred to ice-cold slicing solution containing (in mM): 87 NaCl, 26 NaHCO_3_, 2.5 KCl, 10 D-glucose, 1.25 NaH_2_PO_4_, 50 sucrose, 0.5 CaCl_2_, and 3 MgCl_2_. 250 µm thick coronal slices spanning CnF and PAG (-5.2 to -4.1 mm from bregma) were cut using a vibratome (VT 1200S, Leica Biosystems) at a speed of 0.06 mm/s. The slices were subsequently transferred to the same slicing solution, at near-physiological temperature (36 °C) for 30 min before being transferred to artificial cerebrospinal fluid (aCSF; warmed to 36 °C) containing (in mM): 125 NaCl, 2.5 KCl, 26 NaHCO_3_, 1 NaH_2_PO_4_, 10 D-glucose, ∼2 CaCl_2_, and 1 MgCl_2_. The slices were cooled down and stored at room temperature until recordings were obtained. All solutions were equilibrated with carbogen gas (95% oxygen and 5% carbon dioxide). For recordings, slices were transferred to a submerged chamber and continuously perfused with carbogenated aCSF (30–32 °C, ∼2 ml/min perfusion speed). Locations of CnF and PAG neurons in coronal sections of the midbrain were identified using a Slicescope upright microscope with oblique contrast (Scientifica) and a 4x or 40x immersion objective (Olympus). Recording pipettes were prepared using filamented borosilicate glass (Harvard Apparatus, 0.85 mm ID, 1.5 mm OD) and pulled on the day of use with a micropipette puller (Model P-97, Sutter Instrument) to a pipette resistance of 3–5 MΩ. The pipettes were backfilled with internal solution containing (in mM): 130 KMeSO_3_, 10 HEPES, 10 Na-phosphocreatine, 2 Na_2_ATP, 2 MgCl_2_, 0.5 Na-GTP, and 0.2 EGTA. A subset of experiments was performed using a caesium-based internal solution containing (in mM): 120 CsMeSO_3_, 10 HEPES, 10 Na-phosphocreatine, 5 QX-314, 4 Na-ATP, 4 MgCl_2_, 0.3 Na-GTP, 0.5 EGTA, and 0.1 spermine. The internal solution was filtered before use (0.22 µm Millex GV syringe filter, Merck). Whole-cell patch clamp recordings were acquired using a Multiclamp 700B amplifier (Molecular Devices), filtered at 4–10 kHz and digitized with a sampling rate of 25 kHz and 16-bit resolution (Axon Digidata 1550A, Molecular Devices). Optogenetic stimulation was performed using a wide-field LED illuminator (pe-100, CoolLED) at a wavelength of 470 nm and a maximum LED power of 9.14 mW. To test for the presence of light-evoked synaptic inputs, the following protocol was delivered every 10 s: light pulses were delivered with a duration of 1 ms; a single pulse was followed by a pulse train consisting of five pulses at 20 Hz (interpulse interval: 50 ms) with a 2.5 s delay between them. Neurons were recorded in voltage-clamp configuration at a membrane potential of -70 or -80 mV to record excitatory postsynaptic currents (EPSCs) and at -40 or 0 mV to record inhibitory postsynaptic currents (IPSCs); the access resistance was monitored throughout the recording using a brief hyperpolarizing voltage step, and no access resistance compensation was applied. For current clamp recordings, bridge balance correction was performed and, where necessary, a holding current was applied to keep the membrane voltage at -55 to -65 mV.

#### Data analysis

Liquid junction potentials were not corrected. The access resistance was calculated from the current response to a hyperpolarizing voltage step of 5 mV applied per sweep. Only cells with an average access resistance below 35 MΩ were analysed, and sweeps with an access resistance increase of more than 35% were excluded. For the analysis of voltage-clamp recordings, sweeps were low-pass filtered at 5 kHz. The maximum amplitude and synaptic delay of light-evoked synaptic inputs were calculated from the average current response of each cell recorded in voltage-clamp configuration across the five-pulse train after baseline subtraction, at each respective holding potential. If cells exhibited polysynaptic responses, the first peak per pulse was used. The paired pulse ratio across the pulse train was calculated as the ratio of peak amplitudes between the second to fifth and first EPSC. The latency between excitatory and inhibitory synaptic currents was calculated based on their respective synaptic latencies recorded consecutively in the same neuron. The probability of a neuron to fire action potentials in response to optogenetic stimulation was calculated from the same pulse train protocol delivered in current clamp configuration at a membrane potential of -55 to -60 mV. For each pulse, the number of spikes was counted from the onset of light stimulation until the next pulse onset, and the average reported across recorded sweeps for each pulse per neuron. To confirm the locations of the recorded neurons post-hoc, an image was taken of the location of the patch pipette using a 4x objective. The recorded slices were fixed in 4% PFA overnight, immunohistochemically stained, and imaged using an Axio Zoom light microscope (Zeiss). The injection site boundaries were delineated by applying Otsu’s automatic thresholding method (61). Viral spread and pipette tip locations are illustrated in the respective sections of the mouse brain atlas (Figure S5C). Only brains where the spread of the AAV-DIO-ChR2 construct was restricted to the PAG were included.

### Immunohistochemistry

Slices were washed for 10 min in PBS, then incubated in blocking solution (3% normal goat serum, 3% normal donkey serum, 0.3% Triton X-100 in PBS) for 1–2 h. Primary antibodies were applied in blocking solution and incubated for 2–3 days on a shaker at 4 °C. Slices were then washed and incubated with secondary antibodies in blocking solution for 2 h on a shaker at room temperature. Slices were again washed, incubated for 5 min with DAPI in PBS (5 µg/ml, Carl ROTH) and mounted on frosted slides (VWR) with SlowFade Gold mounting medium (Life Technologies), coverslipped and stored at 4 °C until imaging. Primary antibodies used in this study were: chicken anti-GFP (#A10262, Invitrogen), rabbit anti-RFP (#600-401-379, Rockland), rabbit anti-GABA (#GTX125988, GeneTex), rat anti-mCherry (#M11217, Invitrogen). Secondary antibodies used were: goat anti-chicken 488 (#A11039, Invitrogen), donkey anti-rabbit 568 (#A10042, Invitrogen), goat anti-rat 568 (#A11077, Invitrogen), and goat anti-rabbit 647 (#ab150083, abcam).

### General data analysis

Data analysis was performed using custom-written code in Python 3.13 and custom code will be made available on request. Data were tested for normal distribution, and reported as mean ± s.e.m. if they passed the normality test or as median (range) if the data were not normally distributed (D’Agostino & Pearson test, alpha = 0.05). Statistical comparisons using significance tests were made using SciPy Stats, and statistical significance was considered when *p* < 0.05. Non-parametric tests were used unless otherwise stated. All post-hoc comparisons were corrected for multiple comparisons. Electrophysiological data were partially analysed in Igor Pro 9 (WaveMetrics, USA) using the software package NeuroMatic (62), and statistical testing performed in Prism 11 (GraphPad).

## Supplementary Figures

**Figure S1.**
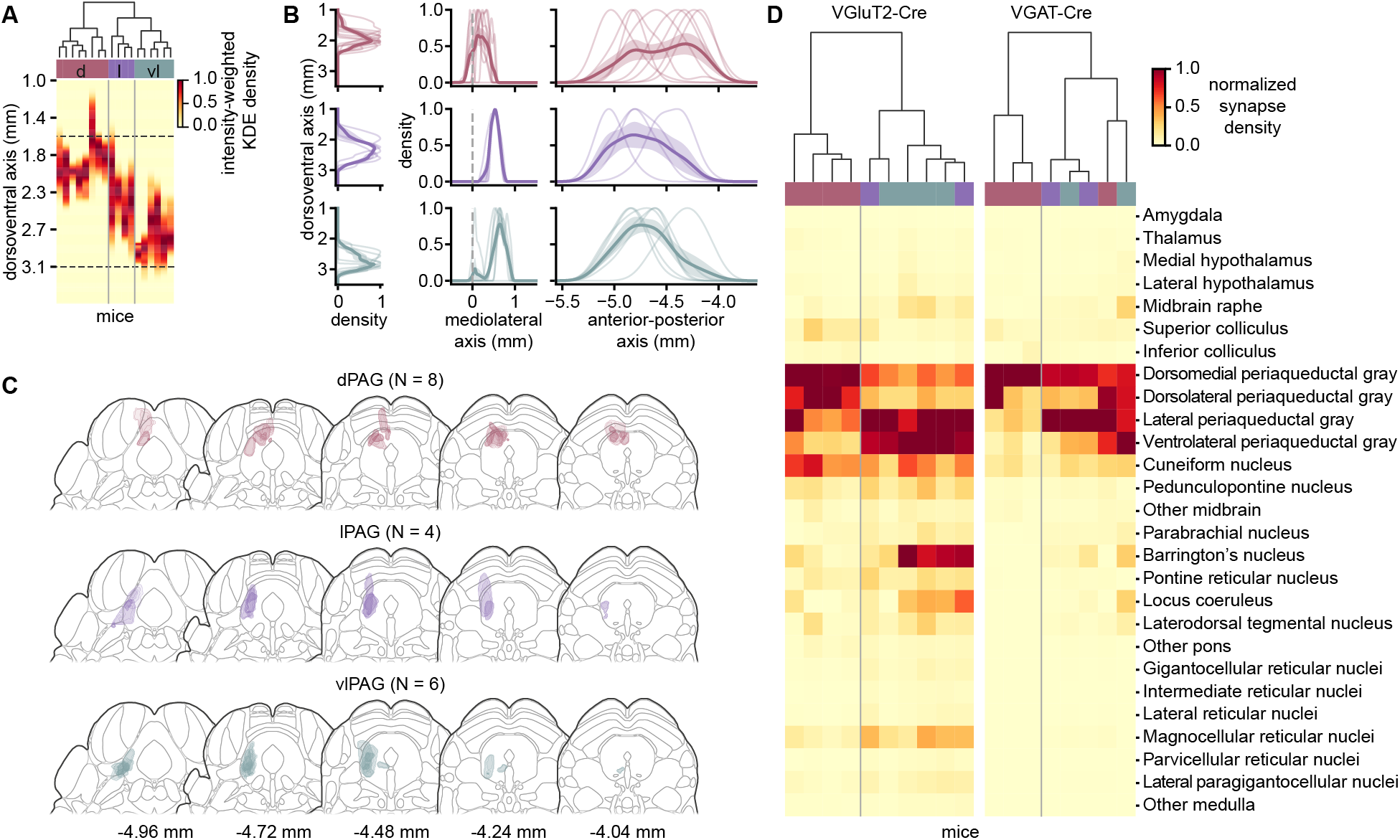
Expression of mGFP in the PAG and synaptic distribution pattern. **(A)** Hierarchical clustering of injection sites based on the intensity-weighted KDE of mGFP fluorescence across the dorsoventral axis for the data in Figure 1 and Figure 2. Dashed lines indicate PAG boundaries. **(B)** 2D KDE of mGFP fluorescence intensity across the dorsoventral (left panels), mediolateral (middle panels), and anterior-posterior (right panels) axis for injection sites clustered into the dorsal (red, top panels), lateral (purple, middle panels), and ventrolateral (teal, bottom panels) part of the PAG. Bold lines show the group mean; faint lines show individual animals. Dashed lines at ML = 0 in mediolateral plots indicate the PAG midline. **(C)** Intensity-thresholded expression of mGFP across all injected animals in the dorsal (red, top panels), lateral (purple, middle panels), and ventrolateral (teal, bottom panels) part of the PAG. **(D)** Hierarchical clustering of regional distribution of synapses from PAG neurons. Each column represents one animal from Figure 1 (VGluT2-Cre, left) or Figure 2 (VGAT-Cre, right); column colour bars indicate PAG subdivision assignment from hierarchical clustering in (A). Shading in (B) is mean ± s.e.m.

**Figure S2.**
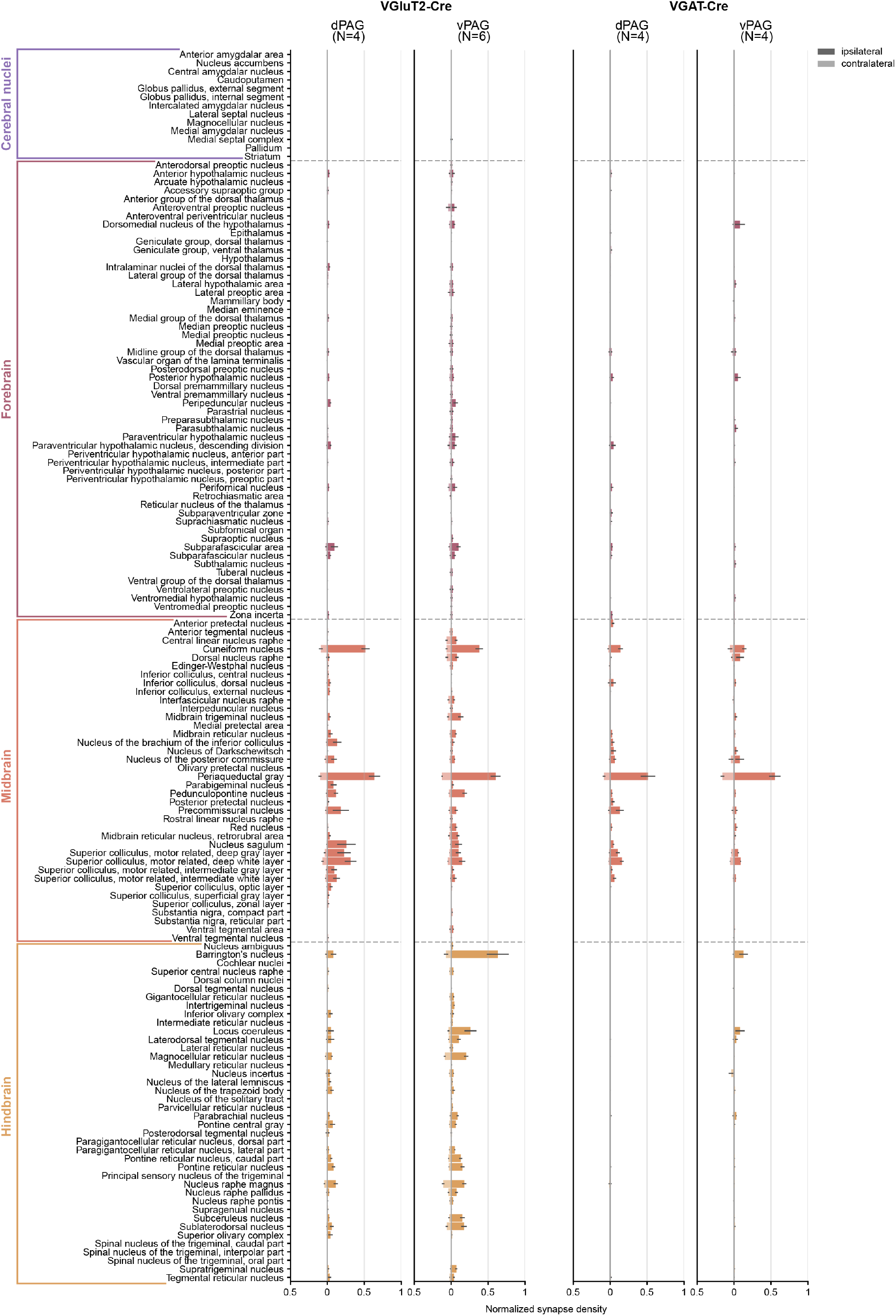
Extended overview of the distribution of synaptic terminals from PAG neurons. Extended region-by-region breakdown of the data summarized in Figures 1G and 2C. Regional density of synapses from VGluT2^+^ (left panels) and VGAT^+^ (right panels) neurons of the dPAG and vPAG, normalized to the densest region per brain. Dark and light bars represent ipsilateral and contralateral hemispheres, respectively. Brain regions are grouped by major division (cerebral nuclei, forebrain, midbrain, hindbrain) based on the Allen CCFv3. Structures are included based on detectable innervation above background or relevance to PAG-associated behavioural systems. Not all analysed structures are included. Error bars are s.e.m.

**Figure S3.**
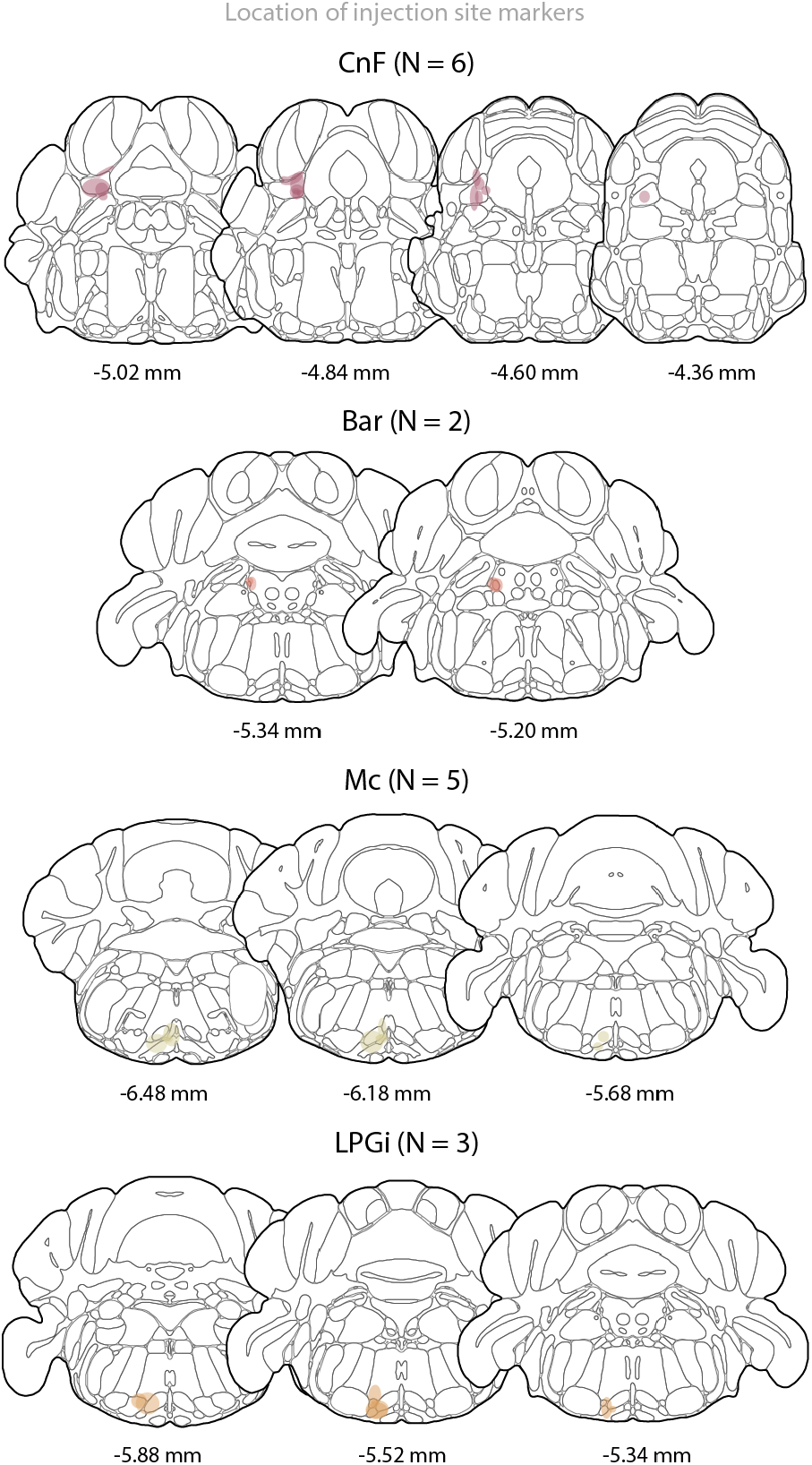
Injection site placement for retrograde labelling. Outlines of expression of the injection site marker enhanced blue fluorescent protein (EBFP) per target region for retrograde labelling shown in Figure 3.

**Figure S4.**
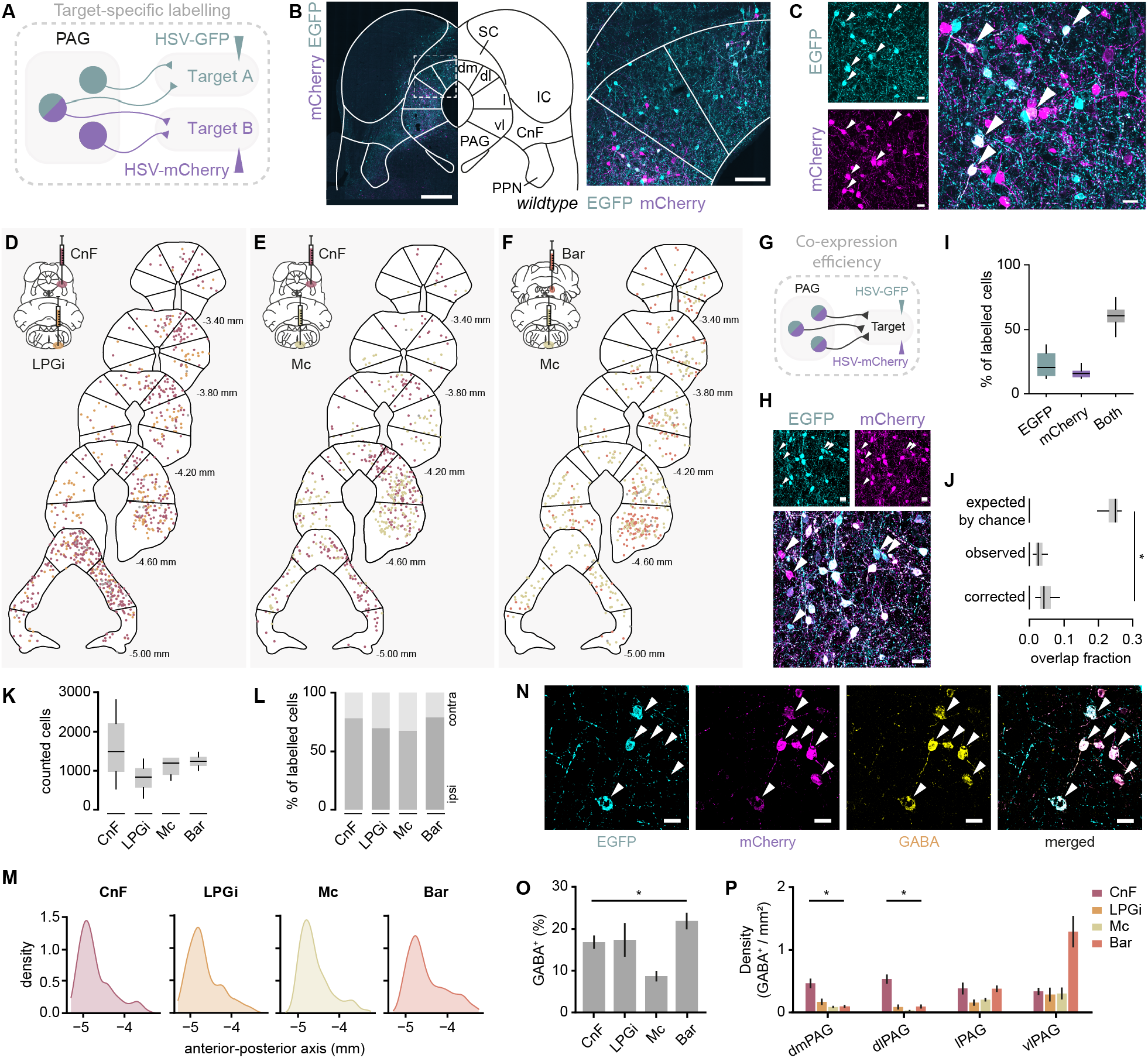
Quantification of viral co-expression efficiency for validation of retrograde labelling strategy, and characterization of GABAergic identity of labelled neurons. **(A)** Target-specific dual-colour retrograde labelling strategy underlying the data in Figure 3. **(B)** Expression pattern of EGFP (cyan) and mCherry (magenta) in PAG neurons of a C57BL/6J wildtype mouse (scale bars: 500 µm left, 100 µm right). **(C)** Expression of EGFP (cyan) and mCherry (magenta) in PAG neurons (scale bars, 20 µm). White arrows indicate cells co-expressing both viral constructs. **(D-F)** Example distribution of retrogradely labelled cell bodies in the PAG along the anterior-posterior axis, downsampled by a factor of 5, from **(D)** CnF (bordeaux) and LPGi (orange), **(E)** CnF (bordeaux) and Mc (ochre), and **(F)** Bar (salmon) and Mc (ochre). **(G)** Strategy for determining co-expression efficiency of viral constructs. **(H)** Co-expression of viral constructs injected into the same brain region (scale bars, 20 µm). White arrows indicate cells expressing only one viral construct, illustrating incomplete co-expression efficiency. **(I)** Percentage of all labelled cell bodies in the PAG expressing EGFP (cyan), mCherry (magenta), or both (grey) after injecting both viral constructs into the same brain region (*N =* 7 mice). **(J)** Comparison of the expected overlap fraction assuming independence of the populations, the observed overlap fraction, and the overlap corrected for the inherent co-expression efficiency of both viral vectors determined in (I). The observed and corrected overlap are significantly smaller than the overlap expected by chance (*N =* 8 mice, paired Wilcoxon signed-rank test: *p* < 0.05 for expected vs. actual and expected vs. corrected). **(K)** Raw stereological counts underlying the estimated totals in Figure 3J. The number of counted cell bodies in the PAG did not differ significantly between projection targets (Kruskal-Wallis test: *p* = 0.511). **(L)** Percentage of retrogradely labelled cell bodies in the PAG located in the ipsi-vs. contralateral hemisphere to the injection site by projection target. **(M)** 2D KDE of the number of retrogradely labelled PAG cell bodies along the anterior-posterior axis by projection target. **(N)** Expression of EGFP (cyan), mCherry (magenta), and anti-GABA (yellow) for cell type determination (scale bars, 20 µm). White arrows indicate GABA^+^ neurons. **(O)** Percentage of GABA^+^ neurons out of all retrogradely labelled cell bodies in the PAG by projection target. The percentage of GABA^+^ neurons was significantly smaller for Mc-projecting PAG neurons (*n* = 88 slices in *N =* 6 mice, linear mixed model (Wald F): *p* < 0.05, t-test post-hoc comparisons). **(P)** Density of GABA^+^ neurons per PAG column by projection target (linear mixed model (Wald F): *p* < 0.05 (dmPAG and dlPAG), *p* = 0.134 (lPAG), *p* = 0.063 (vlPAG); t-test post-hoc comparisons). Error bars are s.e.m.; box-and-whisker plots show median, IQR, and range; * *p* < 0.05.

**Figure S5.**
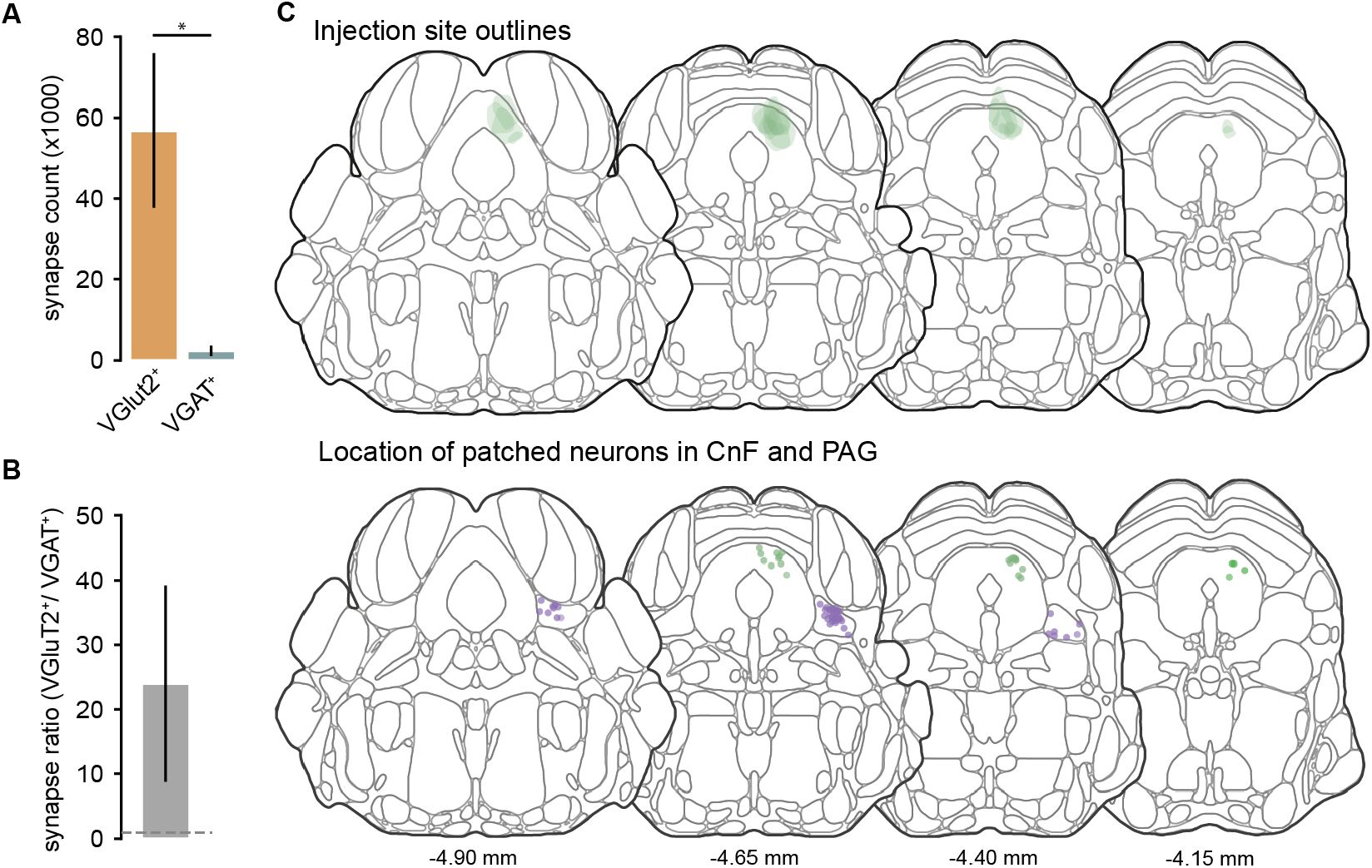
Quantification of dPAG synaptic terminals in CnF, and location of injection sites and recorded neurons for ChR2-assisted circuit mapping. **(A)** Absolute number of presynaptic terminals from VGluT2^+^ (orange) and VGAT^+^ (teal) dPAG neurons detected in the CnF. The number of counted presynaptic terminals was significantly larger for VGluT2^+^ than for VGAT^+^ dPAG neurons (*N =* 8 mice, *p* < 0.05, Mann-Whitney U test). **(B)** Ratio of VGluT2^+^ over VGAT^+^ presynaptic terminals of dPAG neurons in the CnF. The number of VGluT2^+^ synaptic terminals exceeded the number of VGAT^+^ presynaptic terminals by a factor of 24 ± 15. **(C)** Outlines of injection sites for ChR2-expression in dPAG neurons (top panel) and location of individual recorded neurons in dPAG and CnF (bottom panel) for data shown in Figure 4 (*N =* 8 mice). Error bars are s.e.m.; * *p* < 0.05.

## References

1. Bandler, R., Carrive, P., and Depaulis, A. Emerging principles of organization of the midbrain periaqueductal gray matter. In Depaulis, A. and Bandler, R., editors, The Midbrain Periaqueductal Gray Matter: Functional, Anatomical, and Neurochemical Organization, volume 213 of NATO ASI Series A: Life Sciences, pages 1–8. Plenum Press, New York, 1991.

2. Bandler, R. and Shipley, M.T. Columnar organization in the midbrain periaqueductal gray: modules for emotional expression? Trends in Neurosciences, 17(9):379–389, 1994.

3. Behbehani, M.M. Functional characteristics of the midbrain periaqueductal gray. Progress in Neurobiology, 46(6):575–605, 1995.

4. Carrive, P. The periaqueductal gray and defensive behavior: Functional representation and neuronal organization. Behavioural Brain Research, 58(1–2):27–47, 1993.

5. Linnman, C., Moulton, E.A., Barmettler, G., Becerra, L., and Borsook, D. Neuroimaging of the periaqueductal gray: State of the field. NeuroImage, 60(1):505–522, 2012.

6. Stempel, A.V. A conserved brainstem region for instinctive behaviour control: The vertebrate periaqueductal gray. Current Opinion in Neurobiology, 86:102878, 2024.

7. Silva, C. and McNaughton, N. Are periaqueductal gray and dorsal raphe the foundation of appetitive and aversive control? a comprehensive review. Progress in Neurobiology, 177: 33–72, 2019.

8. Bandler, R. and Keay, K.A. Chapter 17 columnar organization in the midbrain periaqueductal gray and the integration of emotional expression. Progress in Brain Research, 107:285–300, 1996.

9. Fanselow, M.S. The midbrain periaqueductal gray as a coordinator of action in response to fear and anxiety. In Depaulis, A. and Bandler, R., editors, The Midbrain Periaqueductal Gray Matter: Functional,Anatomical, and Neurochemical Organization, volume 213 of NATO ASI Series A: Life Sciences, pages 151–173. Plenum Press, New York, 1991.

10. Schenberg, L.C., Bittencourt, A.S., Sudré, E.C.M., and Vargas, L.C. Modeling panic attacks. Neuroscience & Biobehavioral Reviews, 25(7):647–659, 2001.

11. Depaulis, A., Keay, K.A., and Bandler, R. Longitudinal neuronal organization of defensive reactions in the midbrain periaqueductal gray region of the rat. Experimental Brain Research, 90(2):307–318, 1992.

12. Evans, D.A., Stempel, A.V., Vale, R., Ruehle, S., Lefler, Y., and Branco, T. A synaptic threshold mechanism for computing escape decisions. Nature, 558(7711):590–594, 2018.

13. Hoy, J.L., Yavorska, I., Wehr, M., and Niell, C.M. Vision drives accurate approach behavior during prey capture in laboratory mice. Current Biology, 26(22):3046–3052, 2016.

14. Kohl, J., Babayan, B.M., Rubinstein, N.D., Autry, A.E., Marin-Rodriguez, B., Kapoor, V., Miyamishi, K., Zweifel, L.S., Luo, L., Uchida, N., and Dulac, C. Functional circuit architecture underlying parental behaviour. Nature, 556(7701):326–331, 2018.

15. Signoret-Genest, J., Schukraft, N., Reis, S.L., Segebarth, D., Deisseroth, K., and Tovote, P. Integrated cardio-behavioral responses to threat define defensive states. Nature Neuroscience, 26(3):447–457, 2023.

16. Tovote, P., Esposito, M.S., Botta, P., Chaudun, F., Fadok, J.P., Markovic, M., Wolff, S.B.E., Ramakrishnan, C., Fenno, L., Deisseroth, K., Herry, C., Arber, S., and Lüthi, A. Midbrain circuits for defensive behaviour. Nature, 534(7606):206–212, 2016.

17. Tovote, P., Fadok, J.P., and Lüthi, A. Neuronal circuits for fear and anxiety. Nature Reviews Neuroscience, 16(6):317–331, 2015.

18. Verstegen, A.M.J., Klymko, N., Zhu, L., Mathai, J.C., Kobayashi, R., Venner, A., Ross, R.A., VanderHorst, V.G., Arrigoni, E., Geerling, J.C., and Zeidel, M.L. Non-crh glutamatergic neurons in barrington’s nucleus control micturition via glutamatergic afferents from the midbrain and hypothalamus. Current Biology, 29(17):2775–2789.e7, 2019.

19. Vianna, D.M.L. and Brandão, M.L. Anatomical connections of the periaqueductal gray: specific neural substrates for different kinds of fear. Brazilian Journal of Medical and Biological Research, 36(5):557–566, 2003.

20. Zhang, S.P., Davis, P.J., Bandler, R., and Carrive, P. Brain stem integration of vocalization: role of the midbrain periaqueductal gray. Journal of Neurophysiology, 72(3):1337–1356, 1994.

21. Cameron, A.A., Khan, I.A., Westlund, K.N., Cliffer, K.D., and Willis, W.D. The efferent projections of the periaqueductal gray in the rat: A phaseolus vulgaris-leucoagglutinin study. i. ascending projections. Journal of Comparative Neurology, 351(4):568–584, 1995.

22. Cameron, A.A., Khan, I.A., Westlund, K.N., and Willis, W.D. The efferent projections of the periaqueductal gray in the rat: A phaseolus vulgaris-leucoagglutinin study. ii. descending projections. Journal of Comparative Neurology, 351(4):585–601, 1995.

23. Mantyh, P.W. Connections of midbrain periaqueductal gray in the monkey. ii. descending efferent projections. Journal of Neurophysiology, 49(3):582–594, 1983.

24. Tsang, E., Orlandini, C., Sureka, R., Crevenna, A.H., Perlas, E., Prankerd, I., Masferrer, M.E., and Gross, C.T. Induction of flight via midbrain projections to the cuneiform nucleus. PLOS ONE, 18(2):e0281464, 2023.

25. Bouvier, J., Caggiano, V., Leiras, R., Caldeira, V., Bellardita, C., Balueva, K., Fuchs, A., and Kiehn, O. Descending command neurons in the brainstem that halt locomotion. Cell, 163(5): 1191–1203, 2015.

26. Caggiano, V., Leiras, R., Goñi-Erro, H., Masini, D., Bellardita, C., Bouvier, J., Caldeira, V., Fisone, G., and Kiehn, O. Midbrain circuits that set locomotor speed and gait selection. Nature, 553(7689):455–460, 2018.

27. Capelli, P., Pivetta, C., Esposito, M.S., and Arber, S. Locomotor speed control circuits in the caudal brainstem. Nature, 551(7680):373–377, 2017.

28. Esposito, M.S., Capelli, P., and Arber, S. Brainstem nucleus mdv mediates skilled forelimb motor tasks. Nature, 508(7496):351–356, 2014.

29. Falasconi, A., Kanodia, H., Lusk, N., Yao, S., Costa, R.M., Zeng, H., and Arber, S. Projection-defined modules reveal mouse motor cortex architecture. Cell, 2026.

30. Ferreira-Pinto, M.J., Kanodia, H., Falasconi, A., Sigrist, M., Esposito, M.S., and Arber, S. Functional diversity for body actions in the mesencephalic locomotor region. Cell, 184(17): 4564–4578.e18, 2021.

31. Klaus, A., da Silva, J.A., and Costa, R.M. What, if, and when to move: Basal ganglia circuits and self-paced action initiation. Annual Review of Neuroscience, 42(Volume 42, 2019): 459–483, 2019.

32. Mendelsohn, A.I., Nikoobakht, L., Bikoff, J.B., and Costa, R.M. Segregated basal ganglia output pathways correspond to genetically divergent neuronal subclasses. Cell Reports, 44 (4):115454, 2025.

33. Beitz, A.J., Mullett, M.A., and Weiner, L.L. The periaqueductal gray projections to the rat spinal trigeminal, raphe magnus, gigantocellular pars alpha and paragigantocellular nuclei arise from separate neurons. Brain Research, 288(1–2):307–314, 1983.

34. Wang, Q., Ding, S.L., Li, Y., Royall, J., Feng, D., Lesnar, P., Graddis, N., Naeemi, M., Facer, B., Ho, A., Dolbeare, T., Blanchard, B., Dee, N., Wakeman, W., Hirokawa, K.E., Szafer, A., Sunkin, S.M., Oh, S.W., Bernard, A., Phillips, J.W., Hawrylycz, M., Koch, C., Zeng, H., Harris, J.A., and Ng, L. The allen mouse brain common coordinate framework: A 3d reference atlas. Cell, 181(4):936–953.e20, 2020.

35. Mantyh, P.W. Connections of midbrain periaqueductal gray in the monkey. i. ascending efferent projections. Journal of Neurophysiology, 49(3):567–581, 1983.

36. Jansen, A.S.P., Farkas, E., Sams, J.M., and Loewy, A.D. Local connections between the columns of the periaqueductal gray matter: a case for intrinsic neuromodulation. Brain Research, 784(1–2):329–336, 1998.

37. Lau, B.K., Winters, B.L., and Vaughan, C.W. Opioid presynaptic disinhibition of the midbrain periaqueductal grey descending analgesic pathway. British Journal of Pharmacology, 177 (10):2320–2332, 2020.

38. Michael, V., Goffinet, J., Pearson, J., Wang, F., Tschida, K., and Mooney, R. Circuit and synaptic organization of forebrain-to-midbrain pathways that promote and suppress vocalization. eLife, 9:e63493, 2020.

39. Stempel, A.V., Evans, D.A., Arocas, O.P., Claudi, F., Lenzi, S.C., Kutsarova, E., Margrie, T.W., and Branco, T. Tonically active gabaergic neurons in the dorsal periaqueductal gray control instinctive escape in mice. Current Biology, 34(13):3031–3039.e7, 2024.

40. Bindi, R.P., Guimarães, C.C., Oliveira, A.R., Melleu, F.F., Lima, M.A.X., Baldo, M.V.C., Motta, S.C., and Canteras, N.S. Anatomical and functional study of the cuneiform nucleus: A critical site to organize innate defensive behaviors. Annals of the New York Academy of Sciences, 1521(1):79–95, 2023.

41. van der Zouwen, C.I., Tello, A.J., Suresh, J.S., Duque-Yate, J., Hsu, T., Konanur, V., Boutin, J., Roitman, M.F., and Ryczko, D. Cell-type specific sensory and motor activity in the cuneiform nucleus and pedunculopontine nucleus in mice. Scientific Reports, 15(1):20408, 2025.

42. Reichling, D.B. and Basbaum, A.I. Collateralization of periaqueductal gray neurons to forebrain or diencephalon and to the medullary nucleus raphe magnus in the rat. Neuroscience, 42(1):183–200, 1991.

43. Verstegen, A.M.J., Vanderhorst, V., Gray, P.A., Zeidel, M.L., and Geerling, J.C. Barrington’s nucleus: Neuroanatomic landscape of the mouse “pontine micturition center”. Journal of Comparative Neurology, 525(10):2287–2309, 2017.

44. Josset, N., Roussel, M., Lemieux, M., Lafrance-Zoubga, D., Rastqar, A., and Bretzner, F. Distinct contributions of mesencephalic locomotor region nuclei to locomotor control in the freely behaving mouse. Current Biology, 28(6):884–901.e3, 2018.

45. Vaughn, E., Eichhorn, S., Jung, W., Zhuang, X., and Dulac, C. Three-dimensional interrogation of cell types and instinctive behavior in the periaqueductal gray. bioRxiv, page 2022.06.27.497769, 2022.

46. La-Vu, M.Q., Sethi, E., Maesta-Pereira, S., Schuette, P.J., Tobias, B.C., Reis, F.M.C.V., Wang, W., Torossian, A., Bishop, A., Leonard, S.J., Lin, L., Cahill, C.M., and Adhikari, A. Sparse genetically defined neurons refine the canonical role of periaqueductal gray columnar organization. eLife, 11:e77115, 2022.

47. Yeh, L.F., Ozawa, T., Uematsu, A., Duenki, T., Ishizu, Y., Murray, A.J., Okanoya, K., and Johansen, J.P. A sensorimotor brain circuit for transforming aversive experiences into emotional states. bioRxiv, page 2026.01.15.699823, 2026.

48. Streng, M.L., Tetzlaff, M.R., and Krook-Magnuson, E. Distinct fastigial output channels and their impact on temporal lobe seizures. The Journal of Neuroscience, 41(49):10091–10107, 2021.

49. Arber, S. and Costa, R.M. Networking brainstem and basal ganglia circuits for movement. Nature Reviews Neuroscience, 23(6):342–360, 2022.

50. Bizzi, E., Cheung, V.C.K., d’Avella, A., Saltiel, P., and Tresch, M. Combining modules for movement. Brain Research Reviews, 57(1):125–133, 2008.

51. Mussa-Ivaldi, F.A., Giszter, S.F., and Bizzi, E. Linear combinations of primitives in vertebrate motor control. Proceedings of the National Academy of Sciences, 91(16):7534–7538, 1994.

52. Gioia, M., Tredici, G., and Bianchi, R. A golgi study of the periaqueductal gray matter in the cat. neuronal types and their distribution. Experimental Brain Research, 58(2):318–332, 1985.

53. Lefler, Y., Tan, Y.L., Ferreira, G., Fudge, A., Heffernan, M., Wang, Y., and Branco, T. A cellular midbrain mechanism for executing fast and reliable escape. bioRxiv, 2026.

54. Tan, Y.L., Thamilmaran, A., Zernicka-Glover, N., Campagner, D., and Branco, T. Presynaptic temporal dynamics flexibly set input weights in the mouse escape circuit. bioRxiv, 2026.

55. Dean, P., Redgrave, P., and Westby, G.W.M. Event or emergency? two response systems in the mammalian superior colliculus. Trends in Neurosciences, 12(4):137–147, 1989.

56. Dean, P., Redgrave, P., and Mitchell, I.J. Chapter 3: Organisation of efferent projections from superior colliculus to brainstem in rat: evidence for functional output channels. Progress in Brain Research, 75:27–36, 1988.

57. Isa, K., Sooksawate, T., Kobayashi, K., Kobayashi, K., Redgrave, P., and Isa, T. Dissecting the tectal output channels for orienting and defense responses. eNeuro, 7 (5):ENEURO.0271–20.2020, 2020.

58. Roseberry, T., Lee, A., Lalive, A., Wilbrecht, L., Bonci, A., and Kreitzer, A. Cell-type-specific control of brainstem locomotor circuits by basal ganglia. Cell, 164(3):526–537, 2016.

59. Deb, S., Torchia, S., Welponer, E., Spagnoletti, L., Karagülle, S., Aravantis, S., Esteban Masferrer, M., Zhalgasbayev, Y., Asari, H., and Gross, C.T. Reciprocal feedback inhibition and divergent intrinsic firing properties between behaviorally defined classes of neurons in the mouse ventromedial hypothalamus. bioRxiv, 2026.

60. Chiaruttini, N., Castoldi, C., Requie, L.M., Camarena-Delgado, C., Dal Bianco, B., Gräff, J., Seitz, A., and Silva, B.A. Abba+braian, an integrated suite for whole-brain mapping, reveals brain-wide differences in immediate-early genes induction upon learning. Cell Reports, 44 (7):115876, 2025.

61. Otsu, N. A threshold selection method from gray-level histograms. IEEE Transactions on Systems, Man, and Cybernetics, 9(1):62–66, 1979.

62. Rothman, J.S. and Silver, R.A. Neuromatic: An integrated open-source software toolkit for acquisition, analysis and simulation of electrophysiological data. Frontiers in Neuroinformatics, 12:14, 2018.

